# Genetic and chemical validation of *Plasmodium falciparum* aminopeptidase *Pf*A-M17 as a drug target in the hemoglobin digestion pathway

**DOI:** 10.1101/2021.11.23.469631

**Authors:** Rebecca C.S. Edgar, Ghizal Siddiqui, Kathryn Hjerrild, Tess R. Malcolm, Natalie B. Vinh, Chaille T. Webb, Christopher A. MacRaild, Natalie A. Counihan, Darren J. Creek, Peter J Scammells, Sheena McGowan, Tania F. de Koning-Ward

## Abstract

*Plasmodium falciparum,* a causative agent of malaria, continues to remain a global health threat since these parasites have developed increasing resistance to all anti-malaria drugs used throughout the world. Accordingly, drugs with novel modes of action are desperately required to combat malaria. *P. falciparum* parasites infect human red blood cells where they digest the hosts main protein constituent, hemoglobin. Leucine aminopeptidase *Pf*A-M17 is one of several aminopeptidases that have been implicated in the last step of this digestive pathway. Here we utilize both reverse genetics and a compound specifically designed to inhibit the activity of *Pf*A-M17 to show that *Pf*A-M17 is essential for *P. falciparum* survival as it provides parasites with free amino acids for growth, many of which are highly likely to originate from hemoglobin. We further show that our inhibitor is on-target for *Pf*A-M17 and has the ability to kill parasites at nanomolar concentrations. Thus, in contrast to other hemoglobin-degrading proteases that have overlapping redundant functions, we validate *Pf*A-M17 as a potential novel drug target.

## Introduction

Malaria is an infectious disease caused by protozoan parasites belonging to the *Plasmodium* genus, of which *Plasmodium falciparum* is the deadliest to humans. In 2019, there were more than 400,000 deaths attributed to malaria infections, the majority of these occurring throughout sub-Saharan Africa and South-East Asia (1). The current front-line therapeutic artemisinin and its derivatives, as well as partner drugs, are under threat as resistance to these drugs is spreading. Artemisinin resistance is now widespread throughout Asia, and has recently spread to Papua New Guinea, while resistance in malaria endemic regions of Africa emerged last year (2–4). The growing spread of resistance highlights the need to identify new therapeutic targets and compounds with novel modes of action (5).

The intra-erythrocytic cycle of *P. falciparum* is responsible for the clinical manifestations of disease and is the target of most antimalarials. Parasite survival during the erythrocytic stage is dependent upon the digestion of host hemoglobin to provide amino acids essential for parasite growth, with the exception of isoleucine, which is absent from human hemoglobin and, therefore, has to be taken up from the extracellular environment (6). Digestion of hemoglobin also creates space in the erythrocyte to accommodate the growing parasite, as well as providing a mechanism to regulate the osmotic pressure of the host cell (7–10). Hemoglobin digestion begins during the early ring stage of growth, but formation of hemozoin crystals, a detoxified version of the digestive by-product heme, can only be visualized by microscopy in the more developed trophozoite stage in a specialized acidic compartment termed the digestive vacuole (DV; (11). The DV contains an array of proteases responsible for the release of peptides from hemoglobin, including plasmepsins, falcipains, falcilysin and aminopeptidases (12); it is the latter which are speculated to perform the final step of amino acid release from these peptides (13).

As aminopeptidases are implicated in the final step of hemoglobin digestion, they are promising novel therapeutic targets. The *P. falciparum* genome encodes two neutral metallo-aminopeptidases: the M1 alanyl aminopeptidase (*Pf*A-M1) and M17 leucyl aminopeptidase (*Pf*A-M17). The broad-spectrum aminopeptidase inhibitor bestatin has been shown to kill *P. falciparum*, with parasites overexpressing *Pf*A-M17 displaying resistance to this drug, suggesting *Pf*A-M17 is its target *in vivo* (14, 15). Harbut *et al.* (2011) also showed that treatment of *P. falciparum* with bestatin reduced hemoglobin digestion and decreased isoleucine uptake. Specific inhibition of *Pf*A-M17 using an activity-based probe based on the bestatin scaffold resulted in ring-stage arrest and parasite death, whilst an equivalent probe designed to specifically inhibit *Pf*A-M1 resulted in DV swelling and stalling of parasite growth much later at the trophozoite stage. This led the authors to conclude that *Pf*A-M17 may be playing a role outside of, or in addition to, hemoglobin digestion (16). Several series of inhibitors designed to inhibit both *Pf*A-M17 and *Pf*A-M1 have also been developed and these suppress a range of *Plasmodium* species *in vivo* and *in vitro* (17–19). Drinkwater *et al.* (2016), for example, developed a series of dual inhibitors that killed sensitive and multi drug resistant parasites in the nanomolar range, validating *Pf*A-M1 and *Pf*A-M17 as potential novel therapeutic targets.

*Pf*A-M17 is a 68 kDa cytoplasmic enzyme that forms a homo-hexamer in its active form, with optimal function at neutral pH, similar to the pH of the parasite cytoplasm (20–22). Mathew *et al.* (2021) recently confirmed the cytoplasmic localization of *Pf*A-M17, which supports the proposition that hemoglobin-derived peptides are exported from the DV into the parasite cytoplasm, either through the chloroquine-resistance transporter or by other unidentified mechanisms (23). Once in the cytoplasm, it is believed that hemoglobin-derived peptides are digested by *Pf*A-M17, however, it is also possible that *Pf*A-M17 plays an additional role in the catabolic turnover of peptides from other origins. *Pf*A-M17 almost exclusively cleaves leucine and tryptophan *in vitro*, with leucine being one of the most abundant amino acids in adult hemoglobin (24, 25). Functionally, leucine has been shown to be an important substrate of the isoleucine transporter at the parasite membrane (26). As isoleucine is the only amino acid that is absent from hemoglobin and is sourced from the host serum, leucine generated by the parasite may be important for the uptake of isoleucine across the parasite membrane (6, 10). While repeated attempts to knockout *Pf*A-M17 have failed (20, 27), suggesting it is essential for intra-erythrocytic growth, its ortholog could be disrupted in both *Plasmodium berghei*, a rodent malaria species, and in the closely related apicomplexan parasite, *Toxoplasma gondii* (28, 29).

In order to functionally characterize *Pf*A-M17 and determine its contribution to *P. falciparum* survival, we created a conditional knockdown parasite line that enabled *Pf*A-M17 expression to be regulated via a riboswitch (30). This revealed that parasites depleted of *Pf*A-M17 experience a growth delay and fail to expand in culture, subsequently leading to parasite death. We have also designed, synthesized, and characterized a novel and specific small molecule inhibitor to *Pf*A-M17, compound **3**, and parasites treated with this inhibitor demonstrate a similar growth phenotype to *Pf*A-M17 knockdown parasites. Finally, metabolomic analysis of knockdown and **3**-treated parasites revealed that many peptides accumulating after the depletion of *Pf*A-M17 are likely to originate from hemoglobin, indicating that *Pf*A-M17 plays a role in the final stages of its digestion.

## Results

### Epitope tagging and incorporation of a *glmS* ribozyme into the *Pf*A-*m17* locus

To tease out the function of *Pf*A-M17 and determine its essentiality, we sought to use reverse genetics to deplete its expression using a conditional riboswitch system. Accordingly, the *Pf*A-*m17* locus was targeted by transfecting *P. falciparum* 3D7 with a pM17-HAglmS construct (Fig. 1A). Transfectants underwent three rounds of drug cycling with WR99210 before a pure population of integrated parasites was obtained by limiting dilution. Diagnostic PCR confirmed that these parasites were positive for pM17-HAglmS integration (Fig. 1B). Western blot analysis of whole parasite lysate confirmed expression of HA tagged *Pf*A-M17, running slightly lower than the predicted 72 kDa size (Fig. 1C). Immunofluorescence analysis (IFA) further confirmed HA expression, showing that *Pf*A-M17 was excluded from both the food vacuole and the parasite nucleus (Fig. 2A). Sequential solubilization assays performed on mixed-stage parasite lysates showed that *Pf*A-M17 was released into the soluble fraction and was absent from membrane-associated or integral membrane fractions, supporting its cytosolic localization (Fig. 2B). Western blot analysis of protein lysate harvested from *Pf*3D7 wild type parasites every 6 hours and probed with rabbit anti-M17 showed continuous expression throughout the asexual blood stages, with peak expression around 30 hpi as previously reported. (Fig. 2C; See Fig. S1 for M17 antibody characterization).

**Figure 1.**
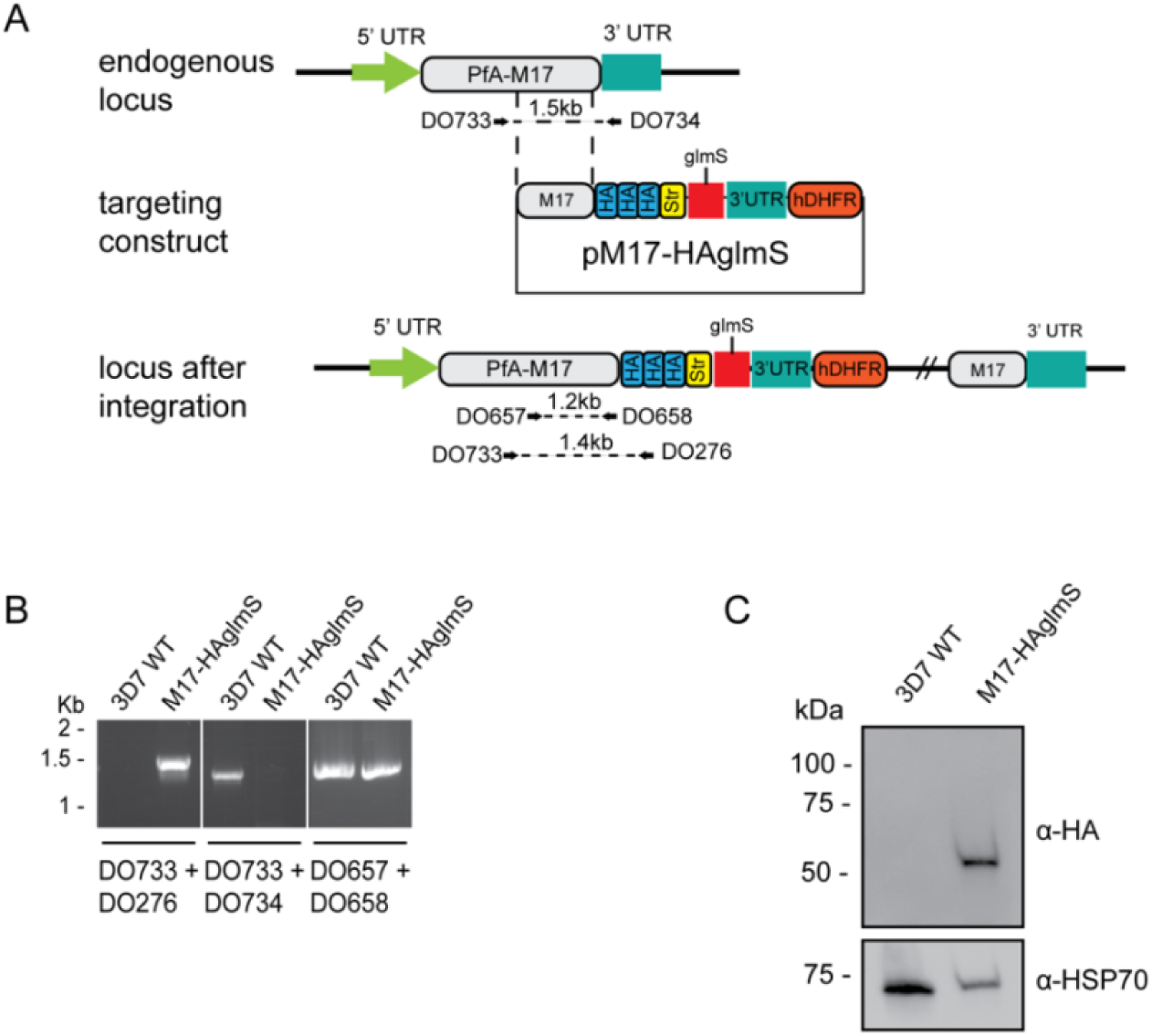
Generation of HA-tagged *Pf*A-M17-HAglmS transgenic parasites. **A**) Schematic of the *Pfa-m17* locus, and locus after single crossover recombination with pM17-HAglmS. The pM17-HAglmS plasmid contained the last kilobase of the coding sequence excluding the stop codon fused in frame to 3 x haemagglutinin (HA) and a single strep II (Str) tag. The plasmid also includes a *glmS* ribozyme with a synthetic untranslated region (UTR) and the selectable marker human dihydrofolate reductase (hDHFR). Arrows indicate oligonucleotides used in diagnostic PCRs as well as their expected sizes. (**B**) Diagnostic PCR showing integration of pM17-HAglmS at the endogenous locus. PCR was performed using the oligonucleotide pairs outlined in (A) on DNA extracted from parasites before (*Pf*3D7) or after (*Pf*A-M17-HAglmS) transfection with the targeting construct. Oligonucleotides DO657 and DO658, which recognize the endogenous locus, serves as a positive control. (**C**) Western blot analysis of parasite lysates confirming HA expression. The predicted molecular mass of *Pf*A-M17-HA is 72 kDa, and HSP70 serves as a loading control.

**Figure 2.**
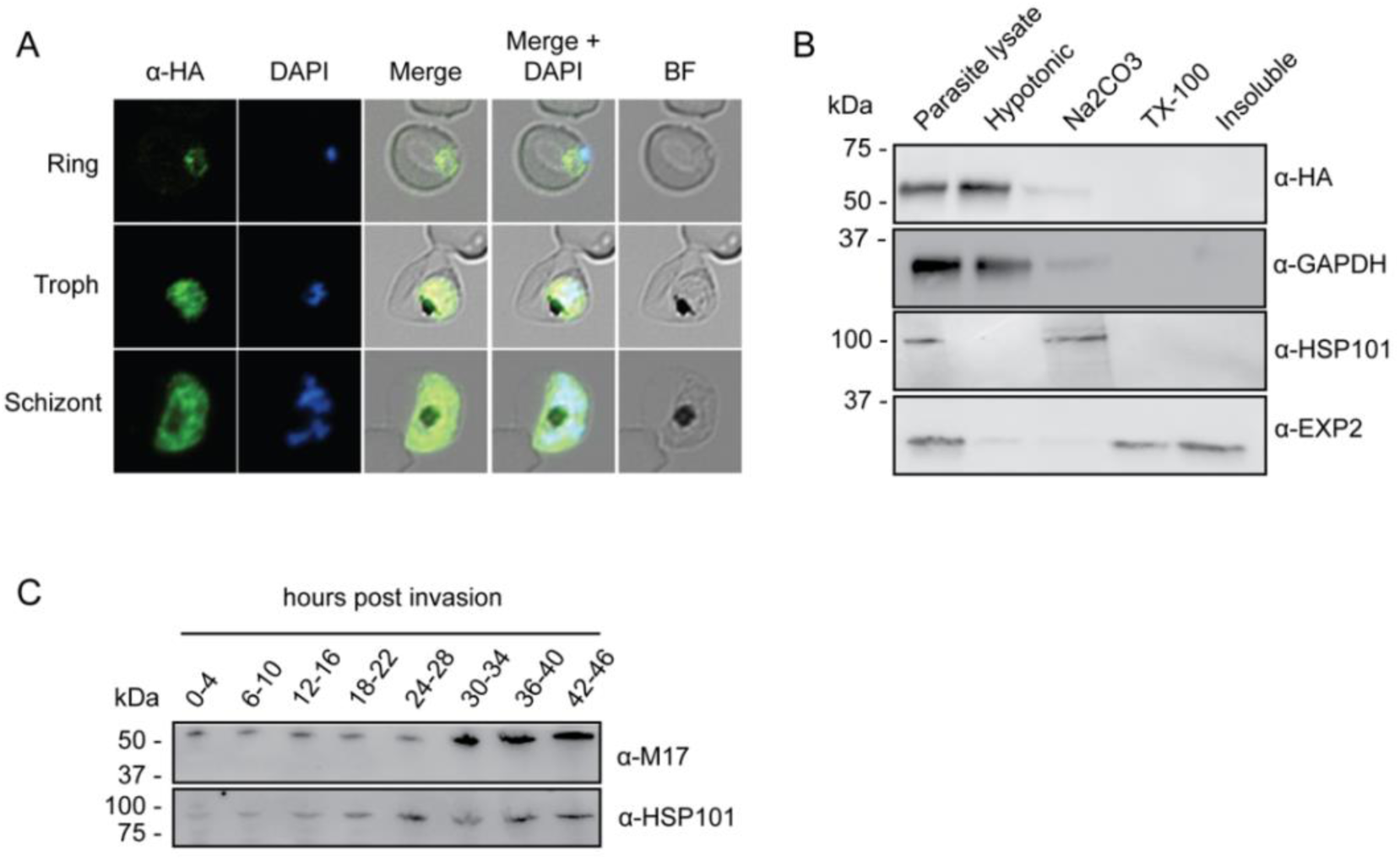
Analysis of *Pf*A-M17 localization and expression over the lifecycle (**A**) Immunofluorescent analysis (IFA) of *Pf*A*-*M17-HAglmS parasites in the three distinct lifecycle stages fixed with 90:10 acetone:methanol and probed with anti-HA and DAPI. (**B**) Saponin-lysed trophozoite stage *Pf*A*-* M17-HAglmS parasites were sequentially lysed in the buffers indicated from left to right and analyzed by Western blotting. Insoluble material represents the remaining pellet after lysis in 1% Triton X-100. GAPDH, HSP101 and EXP2 serve as controls for cytoplasmic, membrane-associated, and integral membrane proteins respectively. (**C**) Western blot analysis of endogenous *Pf*A-M17 expression in *Pf*3D7 wildtype parasites over the erythrocytic cycle probed with anti-M17 antibodies. HSP101 serves as a loading control.

### Knockdown of *Pf*A-M17 expression results in altered parasite growth and a cycle fold-decrease

The synthetic ribozyme incorporated into the 3’UTR of *Pf*A-M17-HAglmS parasites allows the knockdown of protein expression at the transcriptional level with addition of glucosamine (GlcN), allowing characterization of protein function and assessment of the proteins contribution to parasite growth. Ring stage parasites at 0-4 hpi in cycle 1 were treated with 2.5 mM GlcN or left untreated. Parasites were harvested at trophozoite stage in cycle 1 and cycle 2 and significant protein knockdown was determined by Western blotting. This revealed that knockdown was efficient, with 84% and 92% knockdown in cycle 1 and 2, respectively (Fig. 3A). Parasite growth was determined by Giemsa-stained smears and compared to untreated parasites (Fig. 3B). Whilst there was no delay in parasite growth in cycle 1, a significant delay in parasite growth was observed the cycle following knockdown (C2) (Fig. 3B, 3C). This growth delay was already evident by early trophozoites stage and parasites reaching schizogony showed significant morphological changes. Few parasites went on to commence cycle 3, as evidenced by the significantly different parasitemias at 100 h post-treatment (Fig. 3D). Measurement of parasite survival after 10 days in culture was also assessed using a Sybr Green I assay, which revealed knockdown with GlcN was significantly detrimental to parasite growth (Fig. 3E). None of these growth defects were evident in *Pf*3D7 parasites that had been treated with the same concentration of GlcN, confirming that the effect was due to the loss of *Pf*A-M17 (Fig. S2). Overall, this demonstrates that loss of *Pf*A-M17 has a detrimental effect on parasite growth and indicates that this aminopeptidase is essential for survival of *P. falciparum*.

**Figure 3.**
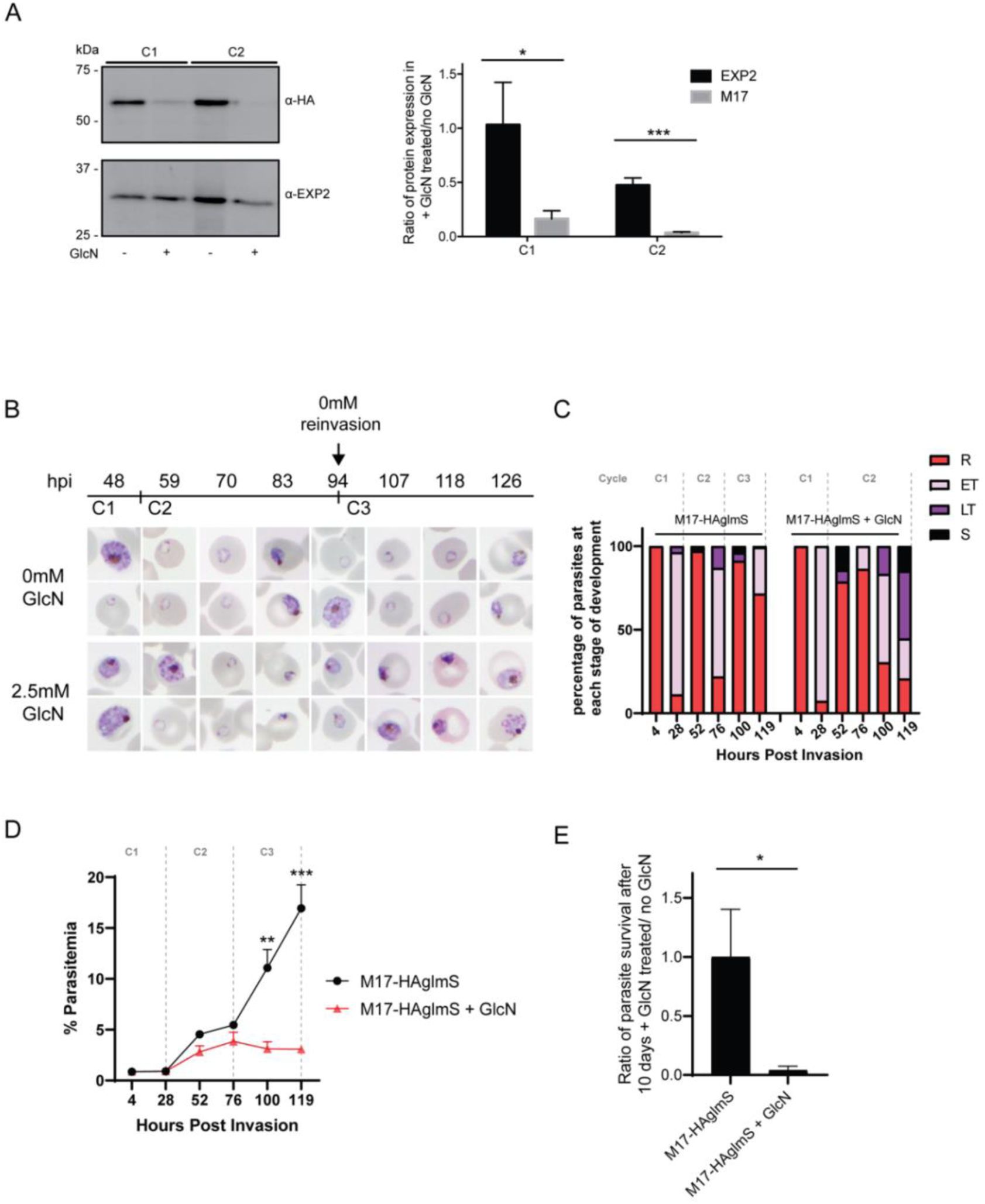
Depletion of *Pf*A-M17 expression leads to perturbed parasite growth *in vitro*. (**A**) Knockdown of *Pf*A-M17 expression. Left panel: Representative Western blot of *Pf*A-M17-HAglmS protein lysates prepared from parasites treated with either 2.5 mM GlcN (+) or left untreated (-) at trophozoite stage in cycle 1 (C1) or cycle 2 (C2). EXP2 serves as a loading control. Right panel: Densitometry of bands observed in Western blots was performed using ImageJ to calculate the ratio of *Pf*A-M17 protein expression in GlcN-treated parasites relative to EXP2 compared to that of untreated parasites. Shown is the mean ± standard deviation (n= 3). Statistical significance was determined using an unpaired t-test, * p ≤ 0.05, *** p ≤ 0.001. (**B**) Representative Giemsa-stained parasite smears of *Pf*A-M17-HAglmS cultures treated with 0 mM or 2.5 mM GlcN shows depletion of *Pf*A-M17 protein results in a growth delay following reinvasion into cycle 2 (C2). (**C**) Percentage of *Pf*A-M17-HAglmS parasites at each stage of development ± GlcN over 3 cycles shows depletion of *Pf*A-M17 leads to delayed parasite development within cycle 2 (n = 3 biological replicates) (**D**) Parasitemias of *Pf*A-M17-HAglmS parasites cultured ± GlcN over 3 cycles. Invasion into cycle 2 is not significantly affected but the growth delay of GlcN-treated parasites observed in this cycle affects parasitemia thereafter, with a significant difference in parasitemia by the time untreated *Pf*A-M17-HAglms have entered cycle 3. Grey dotted lines are representative of the time when untreated parasites have completed a cycle. Plotted is the mean ± standard deviation (n= 3), with statistical significance determined using an unpaired t-test (**p ≤ 0.01, ***p ≤ 0.001). (**E**) Ratio of parasite survival of *Pf*A-M17-HAglmS after treatment with GlcN for 10 days compared to untreated parasites as determined by Sybr Green 1 assay. Shown is the mean ± standard deviation (n= 3). Statistical significance was determined using an unpaired t-test (*p ≤ 0.05).

### Knockdown of *Pf*A-M17 results in multiple digestive vacuoles forming during trophozoite stage but does not alter the quantity of hemozoin

On analysis of *Pf*A-M17-HAglmS parasites the cycle following knockdown, it was noted that some parasites at early trophozoite stage contained multiple digestive vacuoles (DV). To determine the significance of this finding, *Pf*A-M17-HAglmS Giemsa stained parasites, alongside *Pf*3D7 parasites, were scored for their number of digestive vacuoles, indicated by hemozoin under light microscopy, the cycle after GlcN addition; only singly infected RBCs were counted, and the rest excluded (Fig 4A). *Pf*A-M17-HAglmS knockdown parasites had significantly more DV per parasite compared to untreated parasites as determined by a non-parametric Dunn’s post-hoc test (Fig. 4A). The addition of GlcN to *Pf*3D7 parasites did not affect DV numbers, indicating that this phenotype was not as a result of treatment with GlcN but specific to the loss of *Pf*A-M17.

**Figure 4.**
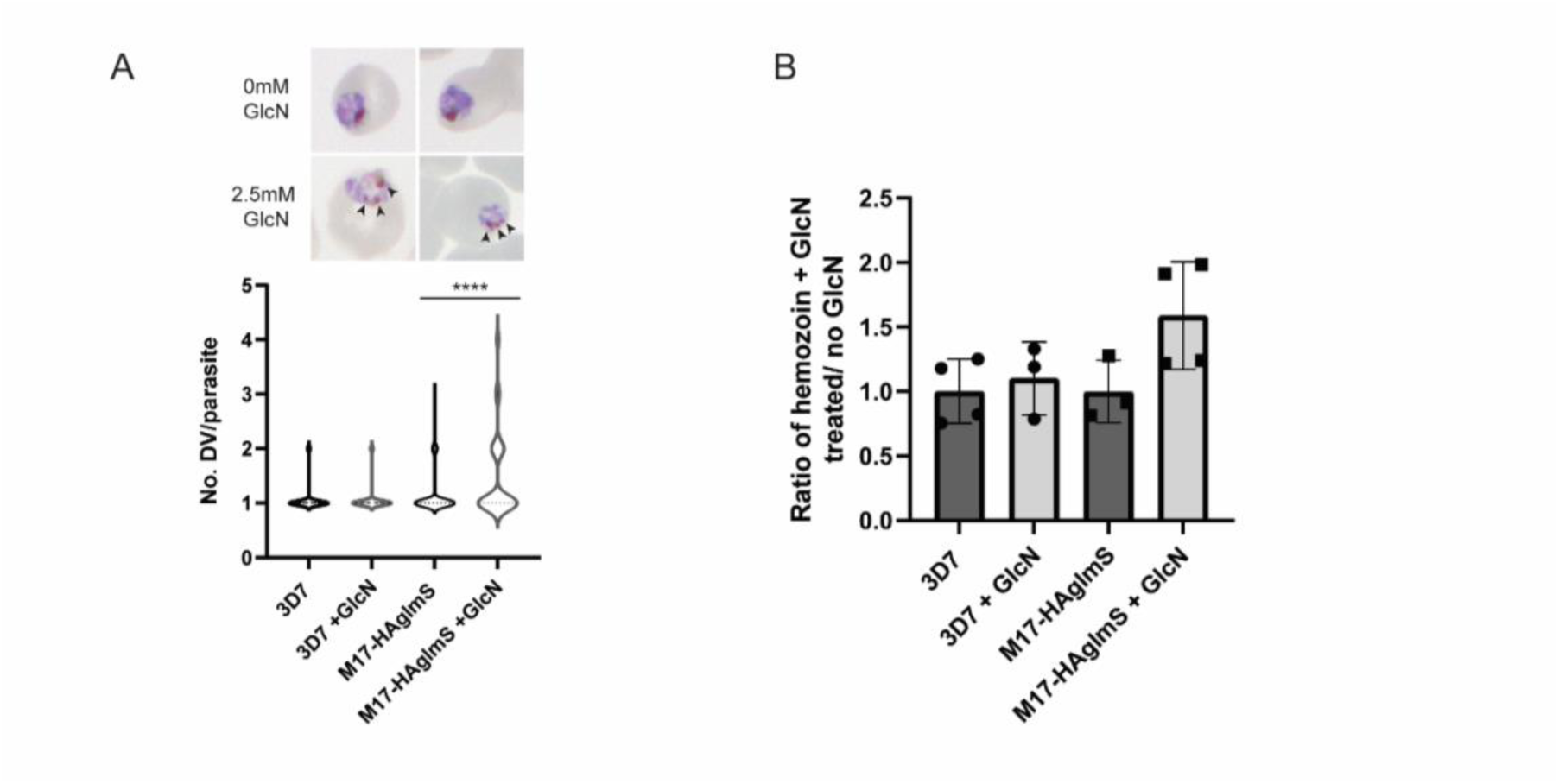
Parasites depleted of *Pf*A-M17 develop significantly more digestive vacuoles the cycle following protein knockdown but hemozoin amount does not change. (**A**) Upper panel: Representative Giemsa-stained smears of *Pf*A-M17-HAglmS ± GlcN; black arrows indicate multiple digestive vacuoles. Lower panel: Number of digestive vacuoles per *Pf*3D7 and *Pf*A-M17-HAglmS the cycle following addition of glucosamine (GlcN) as determined under Giemsa-staining. Shown is combined data from 4 biological replicates (n ≥ 100). Statistical significance was determined by a Kruskal-Wallis test with Dunn’s post hoc multiple comparison test (****p ≤ 0.0001). (**B**) Ratio of hemozoin of *Pf*3D7 and *Pf*A-M17-HAglmS GlcN-treated parasites relative to untreated parasites. Shown is the mean ± standard deviation (n ≥ 3). No statistical significance was determined using an unpaired t-test.

We next determined if the loss of *Pf*A-M17 and increase in number of DV resulted in increased hemozoin formation, and thus hemoglobin digestion. *Pf*A-M17-HAglmS and *Pf*3D7 parasites were harvested at developmentally similar early trophozoite stages the cycle following GlcN addition and the amount of hemozoin was measured. Between all groups there was no statistical difference after the addition of GlcN, and while there appeared to be an upward trend of the amount of hemozoin in GlcN treated *Pf*A-M17-HAglmS parasites compared to untreated, this did not reach significance (Fig. 4B). Thus, while the loss of *Pf*A-M17 increases the number of DV per parasite, it does not increase the quantity of free heme, representative of haemoglobin digestion and hemozoin production, suggesting that there is not an upstream compensatory effect on this digestive pathway.

### A specific inhibitor of *Pf*A-M17 kills parasites within a nanomolar range

Since knockdown of *Pf*A-M17 resulted in parasite death, we next sought to develop and characterize a specific inhibitor to target *Pf*A-M17 and to validate it as on-target using our knockdown parasite line. *Pf*A-M17 uses a metal dependent mechanism to hydrolyze the scissile peptide bond of peptide substrates, removing single amino acids from the N-terminal of short peptides. Our parallel program of inhibitor design and synthesis has identified a number of hydroxamic acid-based compounds that inhibit *Pf*A-M17(19, 31).

A potent and selective *Pf*A-M17 inhibitor, **3** was synthesized from the aryl bromide **1**, that we have reported previously(31). The synthesis first involved the attachment of the 4-hydroxymethylphenyl moiety via a Suzuki coupling reaction to afford **2** (Fig. 5A). The ester of **2** was subsequent converted to the corresponding hydroxamic acid via treatment with hydroxylamine hydrochloride under basic conditions (Fig. 5A). The inhibitory activity toward purified, recombinant *Pf*A-M17 shows **3** to be a potent inhibitor (K_i_ = 18 ± 3 nM) with excellent selectivity over *Pf*A-M1 (K_i_ = 4424 ± 501 nM, Fig.S3), a different aminopeptidase that often shows cross-reactivity with *Pf*A-M17 substrates and inhibitors (19, 31). Analysis of the 2.5 Å X-ray crystal structure of *Pf*A-M17 bound to **3** showed that the position and orientation of the compound was similar to our previous inhibitors with the hydroxamic acid core coordinating to the zinc ions in the active site (Fig. 5B, Fig. S3C). The 4-hydroxymethylphenyl packed with the hydrophobic residues Leu487, Gly489, Leu492 Met396, Phe583 and Ala577 in the S1 pocket and interestingly, the hydroxyl group of **3** appears to interact with the sulfur atom of Met392 (Fig. 5B). The pivalamide occupied the expected position within the S1’ pocket. Further detail and analysis of structure-activity relationships of **3** can be found in Supplementary text.

**Figure 5.**
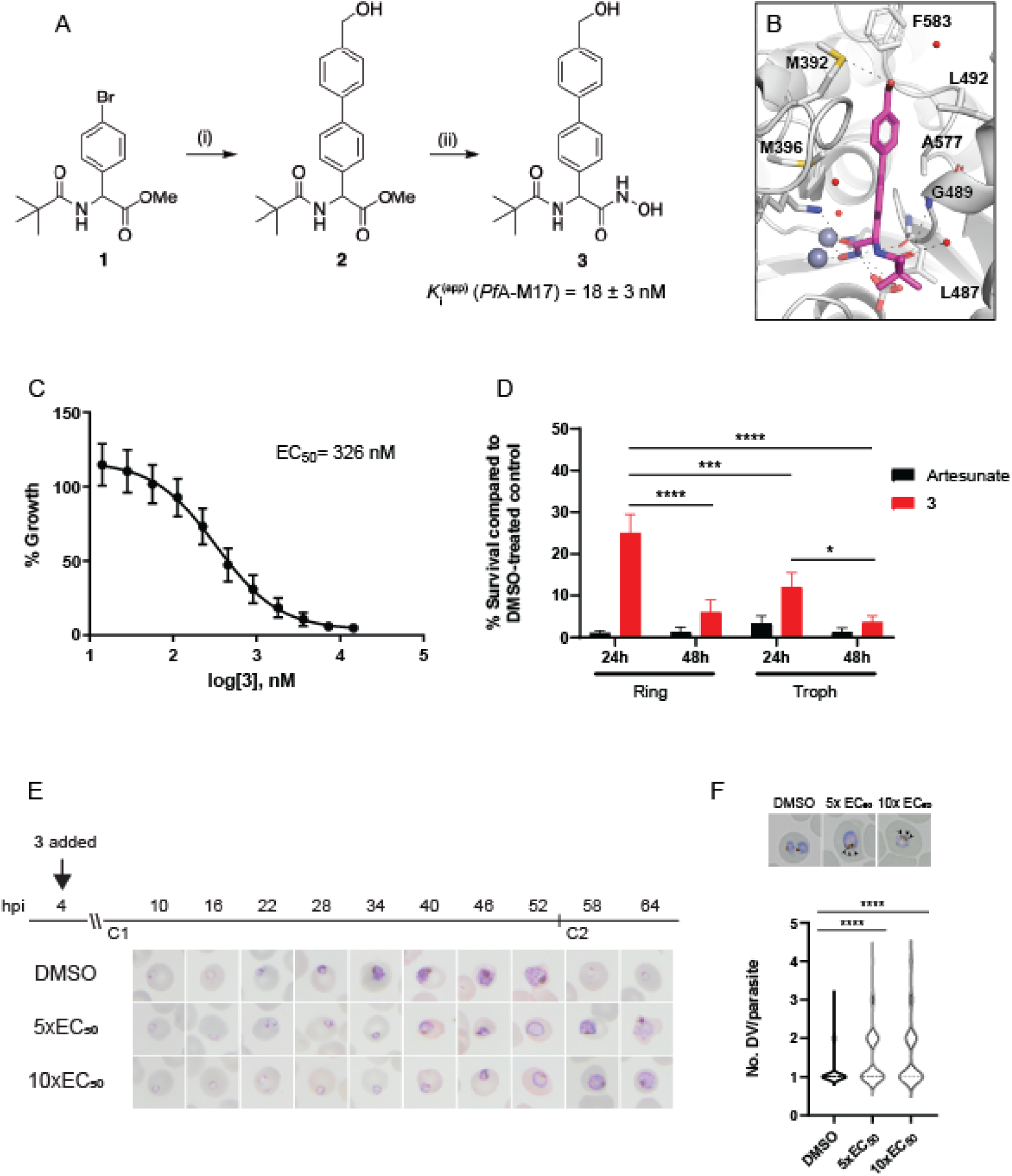
Synthesis and activity of **3,** a specific *Pf*A-M17 inhibitor. **(A)** Scheme 1. Synthesis of **3**: (i) Boronic acid, Pd(PPh_3_)_2_Cl_2_, Na_2_CO_3_, THF, 100 °C, 2 h, (ii) NH_2_OH.HCl, KOH, RT, 16 h. Inhibition constant for **3** toward recombinant, purified *Pf*A-M17 is shown. (**B**) Binding mode of **3** bound to *Pf*A-M17. Solvent-accessible surface of *Pf*A-M17 (grey) with active site ions shown in grey spheres. Stick representation (magenta) shows the binding positions of **3**. Molecular interactions between **3** and *Pf*A-M17 are indicated by dashed lines; water molecules are represented by red spheres. (**C**) Killing action of **3** over 72 h as determined by SYBR Green I assay. The EC_50_ value was calculated from 4 biological replicates performed in triplicate and data plotted as the mean ± standard error of the mean. (**D**) Parasite killing rate was determined by incubating *Pf*3D7 parasites in 10x EC_50_ as previously determined for either 24 or 48 hours before the drug was washed off and parasites allowed to grow for a further 48 hours. Survival was determined via Sybr Green I assay and compared to vehicle (DMSO)-treated controls. Shown is the mean ± standard deviation (n=4) (**E**) Synchronized parasites at 4 hours post-invasion (hpi) were treated over two cycles (C1, cycle 1; C2, cycle 2) with either 5x or 10x EC_50_ or DMSO at the concentration present in the 10x EC_50_ treatment. Representative Giemsa-stained smears from 2 biological replicates show delay in parasite maturation to schizogony (5x EC_50_) or trophozoite stage (10x EC_50_). (**F**) Upper panel: Representative Giemsa-stained smears of *Pf*3D7 + DMSO, 5x or 10x the EC_50_ of **3**; black arrows indicate multiple digestive vacuoles. Lower panel: Number of digestive vacuoles per *Pf*3D7 following the addition of **3** at 4 hours post invasion (hpi) as determined under Giemsa-staining. Shown is the combined data from 2 biological replicates (n ≥ 100). Statistical significance was determined by a Kruskal-Wallis test with Dunn’s post hoc multiple comparison test (****p ≤ 0.0001).

With the inhibitor in hand, its effectiveness on parasites was next tested. The EC_50_ of **3** on *Pf*3D7 was determined to be 326 nM (145-581 CI) using a standard 72-hour killing assay, whereby parasite survival was determined over a range of compound concentrations (Fig. 5C). To determine at what stage of the erythrocytic cycle **3** impacts on parasite growth, parasite cultures were treated with either 10xEC_50_ **3** or artesunate (EC_50_ 4.0 nM, 0.5-6.5 CI) for 24 or 48 hours, commencing at either the ring or trophozoite stage, and following compound washout, cultures were incubated for a further 48 hours (Fig. 5D). This revealed that **3** was less effective at killing parasites with only a 24-hr treatment period compared to the 48-hr treatment, irrespective of whether the compound was administered at ring or trophozoite stage. Treatment for only 24 hours was significantly more effective when added at the trophozoite stage compared to ring stage addition. The most effective killing for **3** was observed when trophozoite stage parasites were treated for 48 hours. These findings were consistent with the observation that expression of *Pf*A-M17 peaks within the rapid trophozoite growth stage, with much lower expression during the ring stage. Moreover, it also corresponds with the growth delay observed at trophozoite stage after knockdown of *Pf*A-M17 protein expression. Analysis of Giemsa-stained *Pf*3D7 parasites treated with 5x or 10x the EC_50_ showed that both treatments resulted in a growth delay commencing around early trophozoite stage when compared to the vehicle control DMSO-treated parasites and parasites had either not entered schizogony (10x EC_50_ treatment) or only just completed schizogony (5 x EC_50_ treatment) 64 hours later (Fig. 5E). Further analysis determined that *Pf*3D7 parasites, upon developing into trophozoites, also developed multiple digestive vacuoles (DV), as seen with specific knockdown of *Pf*A-M17, when treated with either 5x or 10x the EC_50_ of **3** (Fig 5F).

### Analysis of parasites depleted of *Pf*A-M17 alongside treatment with the specific PfA-M17 inhibitor

To determine the specificity of **3** for *Pf*A-M17 and whether it exhibits off-target effects, we next examined the sensitivity of parasites depleted of *Pf*A-M17 to **3**. For these experiments, *Pf*A-M17-HAglmS parasites depleted of *Pf*A-M17 using 2.5 mM GlcN were treated with 5x or 10x the EC_50_ of **3** the cycle following knockdown at the point where the growth delay becomes apparent in GlcN treated parasites (Fig. 3B) and the percentage growth compared to *Pf*A-M17 expressing parasites (ie. not exposed to GlcN treatment) treated with DMSO or 5x the EC_50_. Analysis of growth using Sybr Green I assay showed that there was no significant difference in growth curves between parasites depleted of *Pf*A-M17 and those additionally treated with **3** (Fig. 6A). The reduction in growth at 94 hpi of all treated lines is consistent with the observation that parasites fail to reinvade and commence a subsequent cycle, unlike parasites treated with DMSO alone (Fig. 6A). Moreover, analysis of Giemsa-stained PfA-M17-HAglmS parasites showed that the addition of 5x or 10x the EC_50_ of **3** in the cycle following depletion of *Pf*A-M17 showed a comparative phenotype to *Pf*A-M17 knockdown parasites treated with DMSO (Fig. 6B). These results are in keeping with **3** being on-target and that the effect of this compound on parasites is likely to be through inhibition of *Pf*A-M17.

**Figure 6.**
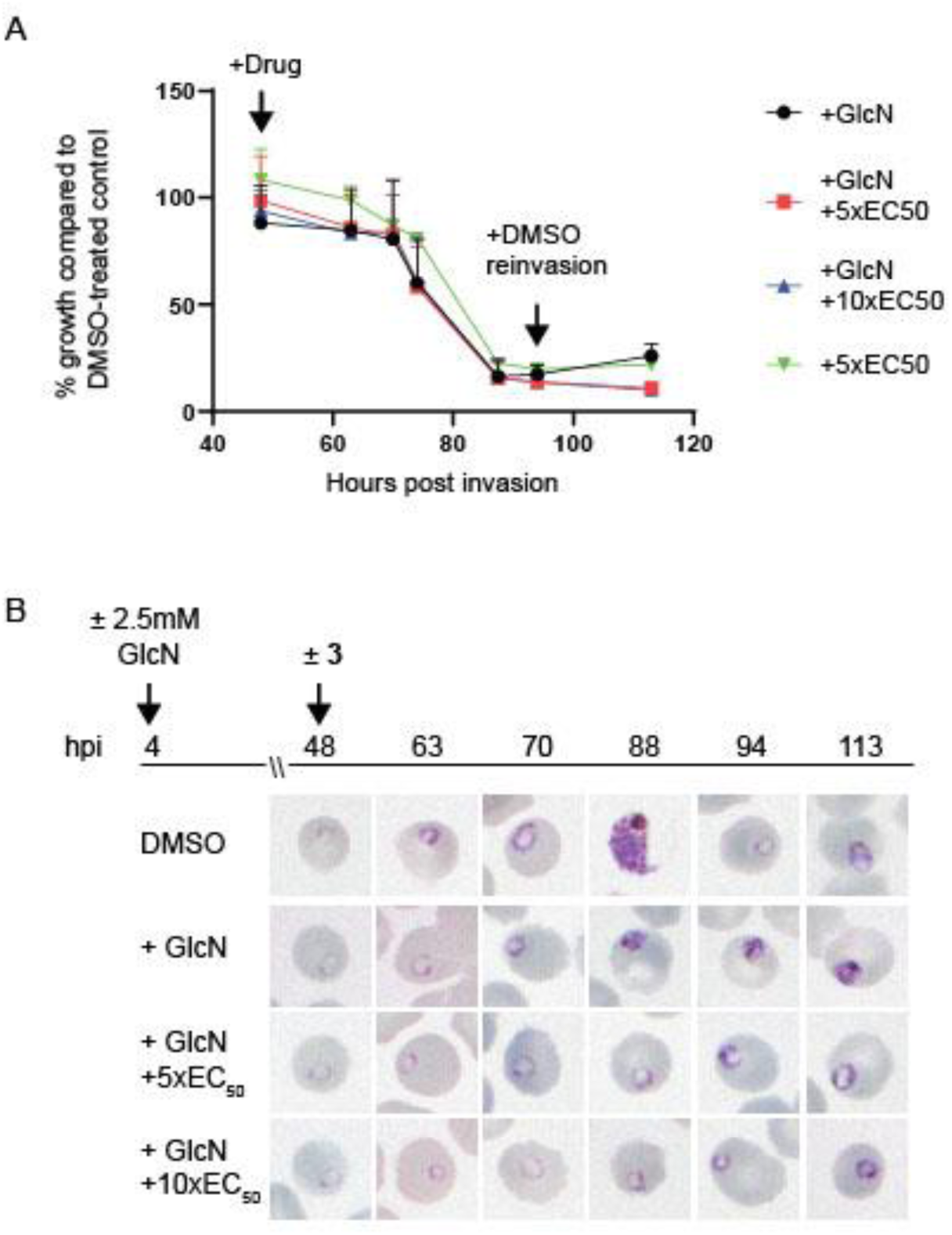
Effect of **3** treatment on parasites depleted of *Pf*A-M17. (**A**) Percentage growth of *Pf*A-M17-HAglmS under different treatment conditions as indicated relative to the growth of DMSO-treated parasites. No significant difference was found between treatments. Plotted is the mean ± standard deviation (n ≥ 2). (**B**) Upper panel: overview of experiment. Heparin synchronized *Pf*A-M17-HAglmS parasites were treated with 2.5mM GlcN and allowed to invade into cycle 2 before being treated with **3** or DMSO. Lower panel: representative Giemsa-stained parasite smears of *Pf*A-M17-HAglmS parasites treated with ± GlcN and ± 5x or 10x EC_50_ of compound **3** shows parasites developing at a slower rate than the DMSO-treated control but in a similar manner between the treatment groups, n=2 biological replicates.

### Metabolomic analysis of parasites depleted of *Pf*A-M17 or treated with 3 indicates *Pf*A-M17 plays a role in the degradation of hemoglobin-derived peptides

To further confirm the specificity of **3** and to determine why knockdown of *Pf*A-M17 has a drastic effect on parasite growth, we compared the metabolomic profiles of *Pf*A-M17 parasites and *Pf*3D7 parasites grown in the presence and absence of GlcN and *Pf*3D7 parasites treated with 10x the EC_50_ of **3** for 1 hour. Principle component analysis and heatmap analysis of relative abundances of putative metabolites dysregulated following *Pf*A-M17 knockdown and parasites treated with **3** showed that the most prominent metabolomic signature shared between them was a series of peptides that were increased (Fig. 7). This was then confirmed by two independent metabolomics experiments of *Pf*A-M17 depletion (Fig. S4), and treatment with **3** (Fig. S5). Targeted analysis of the common set of peptides identified in all three experiments demonstrated that the majority (80 of 149 common peptides) of them are significantly (p-value < 0.05) dysregulated in abundance in parasites depleted of *Pf*A-M17 and following treatment with **3** (Fig. S6). Further detailed analysis of the significantly dysregulated 80 peptides demonstrated that *Pf*A-M17 depletion or treatment with **3** increases the abundance (fold change > 1.5) of these peptides, with the exception of three peptides (Lys/Gly, Glu/Glu/Glu/Lys/Trp, and Asp/Phe/Ile/Tyr/Tyr; indicated by black bar) that were only enriched following treatment with **3** (Fig. 8A). These 80 peptides were then analyzed to determine whether they could be derived from hemoglobin (Hb) α, β, or δ, and ∼82% of these peptides could be mapped to one of the Hb species (yellow dots; Fig. 8B). Thus, metabolomics analysis of parasites depleted of *Pf*A-M17 or treated with a M17 specific inhibitor suggests that *Pf*A-M17 is highly likely to catalyze a terminal stage of Hb digestion.

**Figure 7.**
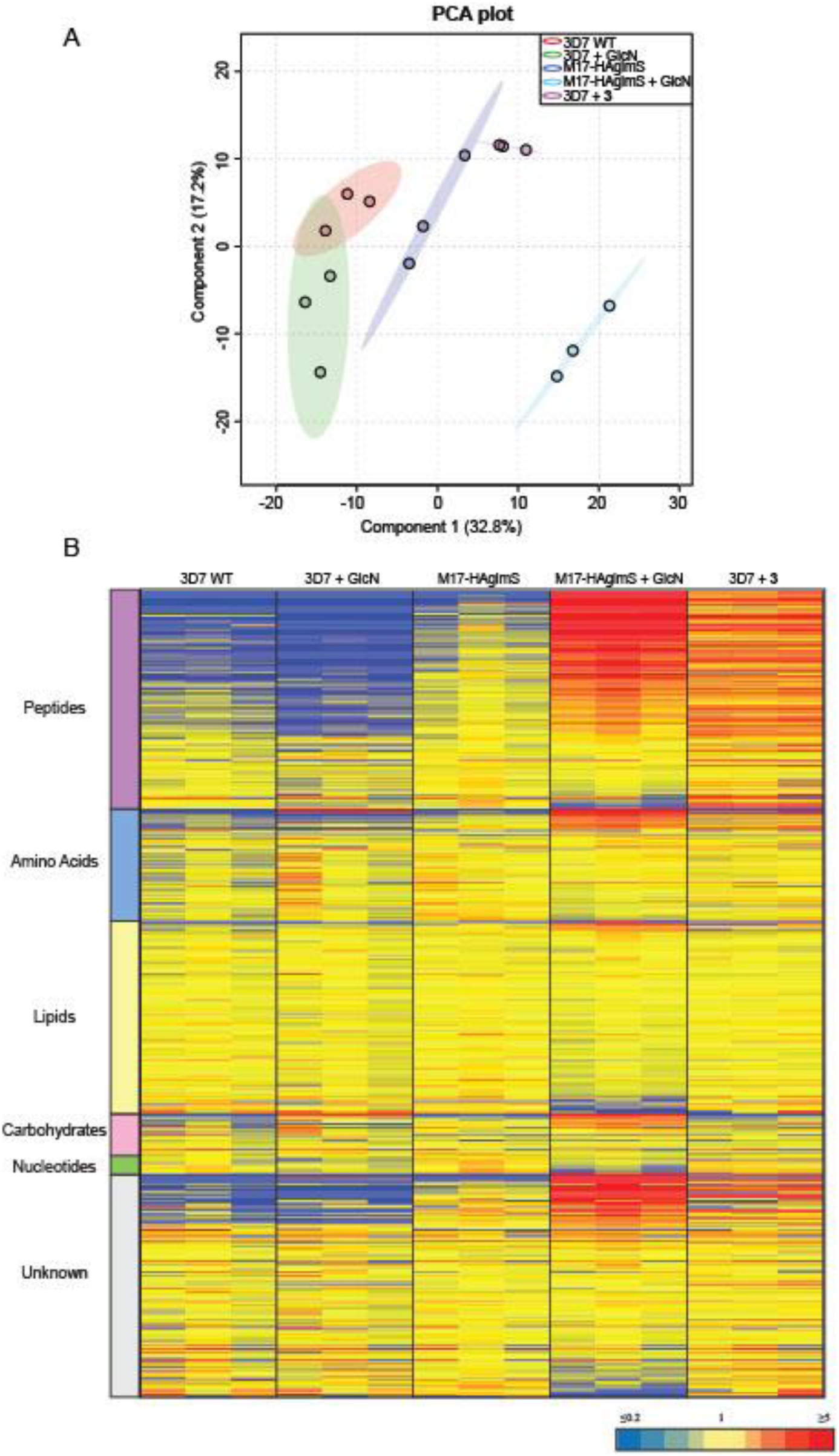
Untargeted metabolomics analysis of *Pf*A-M17-HAglmS and *Pf*3D7 parasites treated with ± GlcN and of *Pf*3D7 parasites treated with **3** from experiment 1. (**A**) Principal component analysis (PCA) of parasites (*Pf*A-M17-HAglmS and *Pf*3D7) treated with +/- GlcN, **3** or DMSO control. Scores plot shows principal components one and two, data points indicate individual sample replicates within each condition and the shaded area denotes 95% confidence interval. (**B**) Heatmap analysis of normalized peak intensities of all putative metabolites for each condition. Data is shown from three technical replicates, red, blue and yellow indicates increase, decrease and no change respectively in the relative abundance of putative metabolites identified.

**Figure 8.**
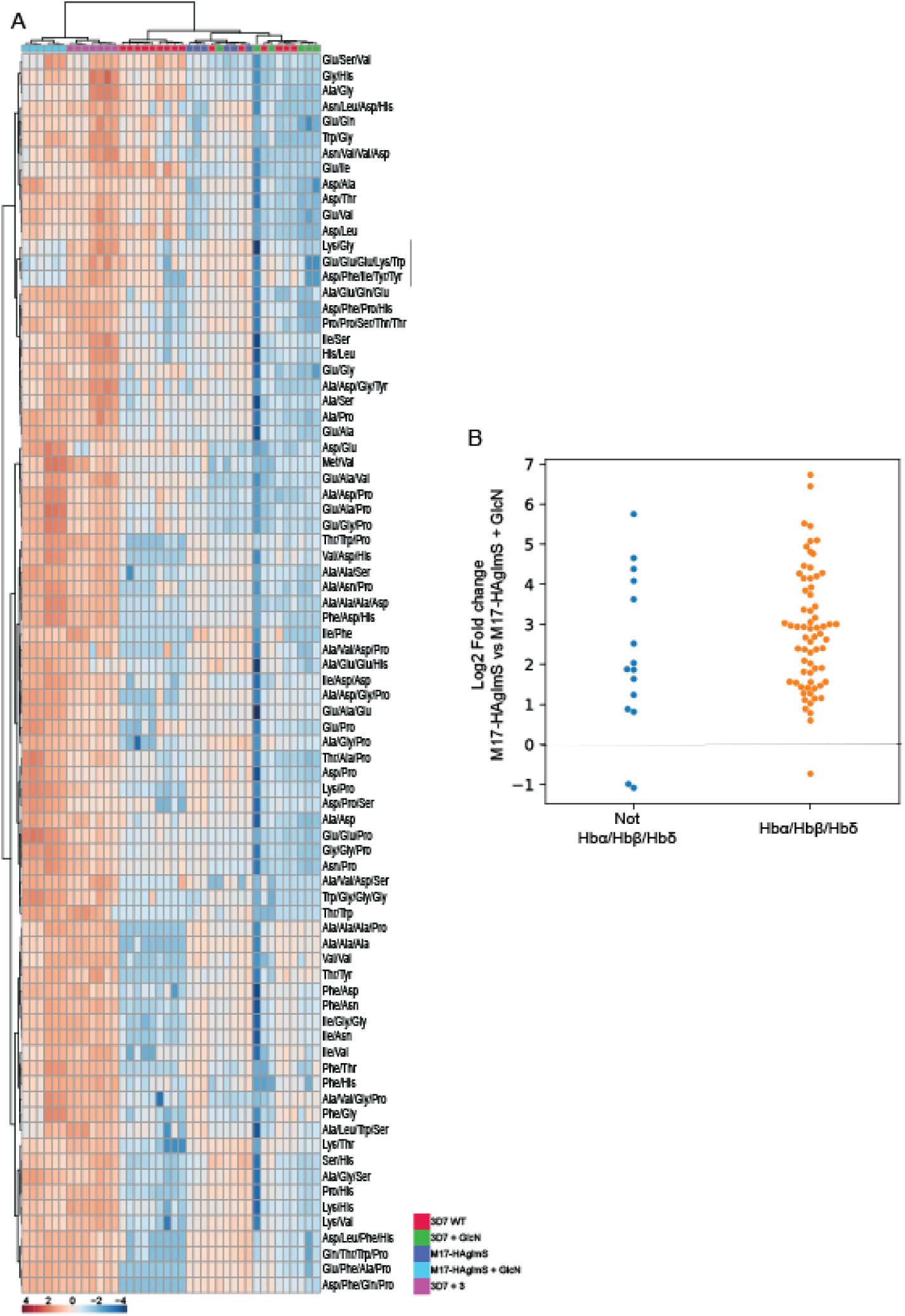
Targeted analysis of common significantly perturbed peptides (p-value < 0.05) following addition of GlcN or treatment with **3** identified from experiment 1, 2 and 3. (A) Hierarchical clustering of the 80 significantly perturbed common peptides (fold change > 1.5 and p-value < 0.05) identified across the three independent experiments. Vertical clustering displays similarities between sample groups, while horizontal clusters reveal the relative abundances (median normalized) of the 80 peptides. The color scale bar represents log_2_ (mean-centered and divided by the standard deviation of each variable) intensity values. (**B**) Differential enrichment of the 80 common significantly perturbed peptides that could (orange dots) or could not (blue dots) be derived from hemoglobin (Hb) α, β, and δ.

### Parasites grown in amino acid free medium containing only isoleucine become sensitized to the *Pf*A-M17 inhibitor 3

Since a function of hemoglobin digestion is to provide free amino acids to the parasite, we next determined if removal of exogenous amino acids, with the exception of isoleucine, from the culture medium would sensitize parasites to **3**. A standard 72-hour killing assay was used to determine the EC_50_ of the compound on *Pf*3D7 parasites cultured concurrently in normal medium containing all amino acids, or amino acid free medium containing only isoleucine. This showed that parasites became significantly more sensitive to **3** in amino acid free media, indicating that its target *Pf*A-M17 is responsible for supplying parasites with amino acids essential for survival (Fig. 9). Comparatively, the loss of exogenous amino acids did not significantly impact the EC_50_ of Artemisinin over 72 h as has previously been shown(16).

**Figure 9.**
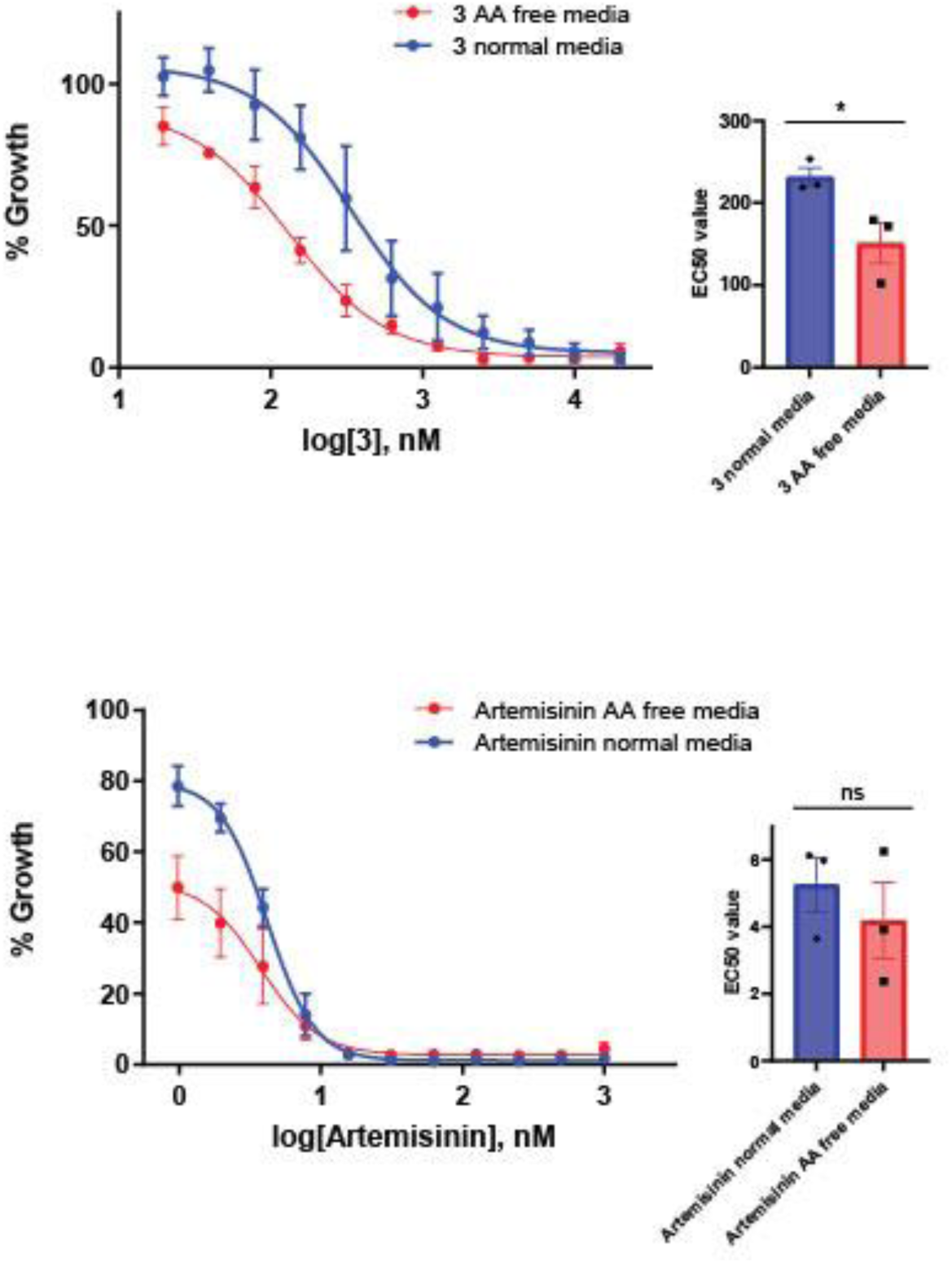
Removal of exogenous amino acids except for isoleucine sensitizes parasites to **3**. Killing action of **3** or Artemisinin in either normal medium containing all amino acids (blue) or amino acid (AA) free medium, containing isoleucine (red) was measured over 72 h and determined by SYBR Green I assay. The EC_50_ values were calculated from 3 biological replicates performed in triplicate and data plotted as the mean ± standard error of the mean, with the inlay bar graphs showing the mean EC_50_ values of these replicates with statistical significance determined using an unpaired t-test, * p ≤ 0.05.

## Discussion

For many years *Pf*A-M17 has been implicated in the final stage of hemoglobin digestion without any definitive confirmation. Here, using a well-established conditional system to knockdown expression of *Pf*A-M17, we demonstrate that a specific loss of this protein corresponds to delayed growth and eventual stalling of parasite development upon transition into trophozoite stage. The resulting phenotype is fatal, with parasites unable to propagate and advance through further cycles, consistent with previous conclusions that *Pf*A-M17 is likely to be an essential protein on the basis that *Pfa-m17* is refractory to gene knockout (20, 27). This validates *Pf*A-M17 as a potential novel drug target.

Unlike previous studies on *Pf*A-M17, which have utilized external activity-based probes or compounds, the conditional knockdown strategy negates any off-target effects that may have convoluted the dissection of its function. Using the approaches herein, we were additionally able to show that **3,** specifically designed to target *Pf*A-M17, was on-target, and parasites treated with this compound displayed a similar phenotype and metabolic profile to parasites in which *Pf*A-M17 expression had been conditionally depleted. Analysis of the significantly disrupted metabolites after *Pf*A-M17 knockdown or treatment with **3** revealed an increase in abundance of peptides, many of which are likely to be derived from hemoglobin. This suggests that the failure to generate a sufficient pool of some amino acids from hemoglobin through a reduction in aminopeptidase activity may be the cause of parasite death. That *P. falciparum* parasites were significantly more sensitized to **3** when cultured in the absence of exogenous amino acids also suggests that the main function of *Pf*A-M17 is to provide amino acids for parasite growth.

Harbut *et al.* (2011) previously showed that *P. falciparum* cultured in the presence of an activity-based probe designed to inhibit *Pf*A-M17 resulted in parasite death at the early ring stage of the parasite lifecycle, suggesting that the role of *Pf*A-M17 in hemoglobin digestion may be subservient to an additional but essential role it plays earlier in the lifecycle (16). The authors hypothesized this additional role is in the turnover of dipeptides originating from the proteosome, as proteasome inhibitors display ring-stage killing (33). While we cannot rule out the latter since some of the peptides that were upregulated upon *Pf*A-M17 knockdown did not map to hemoglobin, this appears not to be the overriding function of *Pf*A-M17 and instead the different stages at which parasites stall in the presence of the activity-based probe are likely to stem from off-target effects.

Teasing out the function of *Pf*A-M17 has been complicated by the fact that another aminopeptidase, *Pf*A-M1, has also been implicated in the final stages of hemoglobin digestion. Initial studies localized *Pf*A-M1 to the digestive vacuole, leading to speculation that the two aminopeptidases play a similar role in different parasite compartments (20). However, it is now clear that *Pf*A-M1 is also cytoplasmic (22). Interestingly, inhibition of *Pf*A-M1 using an affinity-based probe resulted in parasites stalling at the trophozoite stage (16), not dissimilar to the phenotypes observed herein upon knockdown of *Pf*A-M17 or treatment with **3**. However, treatment with this *Pf*A-M1 affinity-based probe also resulted in swelling of the digestive vacuole, a phenotype that we did not observe with *Pf*A-M17 knockdown. This phenotype has previously been seen when enzymes within the digestive vacuole implicated at the earlier stages of hemoglobin digestion have been targeted by compounds (16, 34). Given that *Pf*A-M17 and *Pf*A-M1 are both cytoplasmic proteins, this has begged the question whether there is redundancy between the two, as has previously been shown with other enzymes involved in hemoglobin digestion such as falcipains and plasmepsins (10). Our results, however, indicate that loss of *Pf*A-M17 cannot be compensated for, and as *pfa-m1* has also been previously shown to be impervious to gene knockout, this would suggest that both aminopeptidases are required for parasite survival (20, 27). A thorough analysis of the P1 substrate specificities of recombinant *Pf*A-M17 and *Pf*A-M1 gives some indication as to why this may be. *Pf*A-M1 is a monomeric clan MA alanyl aminopeptidase with broad substrate specificity (25). In contrast, *Pf*A-M17 is a clan MF leucine aminopeptidase with a far narrower specificity for the hydrophobic amino acids, with a strong preference for a P1 leucine and tryptophan residue(25). The inhibitor **3** takes advantage of this specificity with a biphenyl ring system bound in the S1 pocket satisfying the hydrophobic preferences of *Pf*A-M17, but with an addition of a hydroxymethyl group to allow the inhibitor to reach beyond the S1 and pick up an interaction with the S-atom of Met392 (Fig S3C). Taken together, the inhibitor specificity combined with the P1 substrate preference for leucine suggests *Pf*A-M17 may be a dedicated aminopeptidase for processing leucine, with leucine being the most abundant amino acid in human haemoglobin (24).

Since our results indicate that *Pf*A-M17 digests peptides derived from hemoglobin, it is not surprising that this enzyme is not essential in the closely related apicomplexan parasite, *Toxoplasma gondii*, which does not infect red bloods cells and therefore is not exposed to hemoglobin (29). However, the homologous leucine aminopeptidase is also not essential in the rodent malaria species, *P. berghei* (28). Analysis of the *Pb*A-M17 recombinant enzyme identified that it had a similar substrate specificity to that of *PfA*-M17, however, there were notable differences in substrate specificity of the *Pb*-M1 vs *Pf*A-M1 enzymes, suggesting a difference in aminopeptidase substrates between the two species (35). Knockout of *Pb*A-M17 did, however, result in a significant delay in growth, although the significance of this cannot be inferred given the small sample number of two mice (28). Notably, isoleucine is present in mouse hemoglobin but absent in human hemoglobin. In *P. falciparum-*infected red blood cells, the influx of extracellular isoleucine can be mediated by leucine, which serves as a substrate of the isoleucine transporter at the parasite membrane (25, 26). Since *Pf*A-M17 has been demonstrated to have considerable specificity for leucine (25) and *P. berghei* can source isoleucine from hemoglobin, this may explain why *Pf*A-M17 is essential for *P. falciparum* survival whilst *P. berghei* can survive without *Pb*A-M17.

A novel finding observed after specific depletion of *Pf*A-M17 was the presence of multiple digestive vacuoles per parasite, a phenotype not previously seen with inhibition of aminopeptidases in *P. falciparum*. Whether this is because parasites attempt to endocytose more red blood cell cytoplasm to try and salvage their free amino acids pool is unknown and will require further investigation in order to elucidate the mechanism of this finding. We also saw this phenotype in parasites treated with **3**, further suggesting that our compound is on target for *Pf*A-M17. We did not see a significant difference in the amount of hemozoin produced between parasites expressing or depleted of *Pf*A-M17, and a similar finding was observed between *P. berghei* wildtype and *pba-m17* knockout parasites. This indicates that depletion or loss of leucine aminopeptidases may not have an impact further upstream in the hemoglobin digestion pathway resulting in altered hemozoin production.

In conclusion, we have demonstrated that *Pf*A-M17 plays an essential role in the survival of *P. falciparum* and is likely playing a major role in the release of amino acids originating from hemoglobin, confirming *Pf*A-M17 as a promising target for future anti-malarial drugs. Further analysis into additional metabolites found not to map to a hemoglobin sequence will provide insight into their origin and clues as to additional function(s) of *Pf*A-M17. Encouragingly, we have found that **3** is able to kill parasites in a nanomolar range. An X-ray crystal structure of recombinant *Pf*A-M17 shows binding to this compound and addition of **3** to parasites after knockdown of *Pf*A-M17 did not alter the parasite growth or morphology, suggesting the inhibitor is specific for *Pf*A-M17. **3** also appears to be highly selective for *Pf*A-M17 over *Pf*A-M1 and this compound, now validated, provides scope for the further analysis into the function of *Pf*A-M17, negating problems that arise with the death phenotype of the knockdown and providing a rationale for further development of inhibitors against *Pf*A-M17.

## Materials and Methods

### Chemistry

#### Methyl 2-(4’-(hydroxymethyl)-[1,1’-biphenyl]-4-yl)-2-pivalamidoacetate (**2**)

To a mixture of methyl 2-(4-bromophenyl)-2-pivalamidoacetate (400 mg, 1.2 mmol) and 4-(hydroxymethyl)phenylboronic acid (222 mg, 1.5 mmol) in THF (6 mL) was added Na_2_CO_3_ (1 M, 2.0 eq). A steady stream of nitrogen was bubbled through the mixture for 5 min, before PdCl_2_(PPh_3_)_2_ (0.03 eq) was added. The mixture was heated at 100 °C in a sealed tube for 2 h. After cooling, the mixture was diluted with EtOAc (10 mL) and water (10 mL), and the aqueous layer discarded. The organic layer was concentration under reduced pressure and then purified by flash chromatography to give the title compound (385 mg, 89%) of the title compound. ^1^H NMR δ 7.54 (d, *J* = 8.3 Hz, 2H), 7.50 (d, *J* = 8.2 Hz, 2H), 7.40 (d, *J* = 8.3 Hz, 2H), 7.38 (d, *J* = 8.3 Hz, 2H), 6.87 (d, *J* = 6.8 Hz, 1H), 5.57 (d, *J* = 6.8 Hz, 1H), 4.65 (s, 2H), 3.70 (s, 3H), 1.24 (s, 9H); ^13^C NMR δ 178.1, 171.5, 141.0, 140.6, 139.1, 135.3, 127.5, 127.4, 127.3, 126.9, 64.2, 56.1, 52.7, 38.5, 27.2; LC-MS *t*_R_: 3.2 min, *m/z* 356.0 [MH]^+^.

#### N-(2-(Hydroxyamino)-1-(4’-(hydroxymethyl)-[1,1’-biphenyl]-4-yl)-2-oxoethyl)pivalamide (**3**)

Methyl 2-(4’-(hydroxymethyl)-[1,1’-biphenyl]-4-yl)-2-pivalamidoacetate (**2**) (180 mg, 0.51 mmol) and NH_2_OH.HCl (8.0 eq) were dissolved in anhydrous MeOH. KOH (5 M in MeOH, 10 eq) was added and the reaction mixture was stirred at RT overnight. After evaporation of the solvent, the crude product was purified by flash chromatography (eluent MeOH/DCM 0:100 to 10:90) to give the title compound as a white solid (155 mg, 86%). ^1^H NMR (DMSO-*d_6_*) δ 11.00 (s, 1H), 9.05 (s, 1H), 7.70 (d, *J* = 8.0 Hz, 1H), 7.63 (d, *J* = 6.3 Hz, 2H), 7.61 (d, *J* = 6.2 Hz, 2H), 7.48 (d, *J* = 8.3 Hz, 2H), 7.40 (d, *J* = 8.2 Hz, 2H), 5.40 (d, *J* = 8.0 Hz, 1H), 5.20 (t, *J* = 5.7 Hz, 1H), 4.53 (d, *J* = 5.7 Hz, 2H), 1.16 (s, 9H); ^13^C NMR (DMSO-*d_6_*) δ 177.0, 166.9, 141.9, 139.6, 138.3, 138.1, 127.5, 127.2, 126.6, 126.5, 62.8, 53.7, 38.3, 27.3; *m/z* HRMS (TOF ES^+^) C_20_H_25_N_2_O_4_ [MH]^+^ calcd 357.1809; found 357.1818; LC-MS *t*_R_: 3.0 min; HPLC *t*_R_: 5.0 min, > 99%.

### Aminopeptidase activity assays using recombinant purified protein

Recombinant *Pf*A-M17 (amino acids 84-605) and *Pf*A-M1 (amino acids 85-605) expressed in *Escherichia coli* and purified by metal affinity chromatography followed by size-exclusion gel filtration was described previously (19). Aminopeptidase activity was assessed by fluorescence assays using the fluorogenic peptide L-leucine-7-amido-4 methylcoumarin hydrochloride (Sigma L2145) as a substrate as previously described (19). Michaelis−Menten constants (*K*_m_) were calculated for each enzyme purification and showed similar activity as reported previously (21, 36). Inhibition of aminopeptidase activity was measured using a Morrison inhibition constant (K_i_^(app)^), where enzymes were preincubated in 100 mM Tris−HCl, pH 8.0 (supplemented with 2 mM CoCl_2_ for *Pf*A-M17) and compound **3** for 10 min prior to the addition of substrate (20 μM for *Pf*A-M1, 10 μM for *Pf*A-M17). Substrate concentrations did not exceed the *K*_m_ for each enzyme. The inhibitor concentration range was selected to obtain a complete inhibition curve (0−100%). K_i_^(app)^ values were calculated by plotting the initial rates versus inhibitor concentration, and fitting to the Morrison equation for tight-binding inhibitors in GraphPad Prism software (nonlinear regression method). Inhibition constants were calculated in biological triplicate from three different protein preparations. K_i_^(app)^ value represents the mean and standard error of the mean (SEM).

### Structural Biology

*Pf*A-M17 was co-crystallized with **3** by the hanging-drop method, using previously established protocols (19). Briefly, *Pf*A-M17 was concentrated to 10 mg/mL and co-crystallized with a final ligand concentration of 1 mM in 30−40% PEG400, 0.1 M Tris pH 7.5−8.5, 0.2 M Li_2_SO_4_. Crystals were soaked overnight in mother liquor supplemented with 1 mM ligand and 1 mM ZnSO_4_ before being harvested for data collection. Crystals were snap-frozen in liquid nitrogen, and data were collected 100 K using synchrotron radiation at the Australian Synchrotron beamlines 3ID1(MX2). Data was processed using XDS and Aimless as part of the CCP4i program suite. The structures were solved by molecular replacement in Phaser RCSB ID 3KQZ as the search model. The structures were refined using Phenix with 5% of reflections set aside for calculation of R_free_. Between refinement cycles, the protein structure, solvent, and inhibitor were manually built into *2Fo − Fc* and *Fo − Fc* electron density maps using COOT with restraint files generated by Phenix where necessary. Data collection and refinement statistics can be found in Table S1. The coordinates and structure factors are available from the Protein Data Bank with PDB accession code 7RIE.

### Antibody production

Recombinant *Pf*A-M17 (amino acids 84-605;(19)) was used to generate polyclonal rabbit antibodies using The Walter and Eliza Hall Institute of Medical Research Antibody Facility. Briefly, a prebleed was collected from rabbits prior to subcutaneous immunization on four occasions with 200 µg of protein, the first in Freund’s complete adjuvant and the subsequent immunizations in incomplete adjuvant. The final rabbit bleed (and prebleed for comparison) was used to probe for *Pf*A-M17.

### Plasmid constructs

A transgenic *P. falciparum* line allowing conditional knockdown of *Pf*A-M17 was generated by inserting the last 1017 base pairs of *Pf*A-M17 (Pf3D7_1446200), excluding the stop codon, into the *Bgl*II and *Pst*I sites of pPfTEX88-HAglmS using oligonucleotides DO733 and DO734 (37). This resulted in the fusion of the *Pf*A-M17 C-terminus with a triple hemagglutinin (HA) and single streptavidin epitope tag. Oligonucleotide sequences are provided in Table S2.

### Parasite culture and transfection

*P. falciparum* 3D7 and transgenic lines were cultured continuously (38) in O^+^ human erythrocytes obtained from the Australian Red Cross at 4% hematocrit. Complete culturing media contained RPMI 1640 medium (Life Technologies), 20 mg/L gentamicin, 50 mg/L hypoxanthine, 25 mM sodium bicarbonate, 25 mM HEPES and 0.5% (w/v) Albumax II (Life Technologies). Minimal media (ResolvingImages) contained the same constituents as complete media except that isoleucine was the only amino acid present. Parasite cultures were maintained at 37°C at atmospheric conditions of 5% CO_2_ and 1% O_2_ in N_2_. Transfections were performed as previously described (39) and selected for with 2.5 mM WR99210 (Jacobus). Parasite cultures were visualized by making thin blood smears on glass slides, which were fixed with methanol for 10 seconds and then stained with Giemsa (1:10 dilution in water; Merck) for 5-10 min prior to microscopy under 100 x magnification with oil.

### Analysis of *Pf*A-M17 expression over the erythrocytic lifecycle and Western blotting

Erythrocytes infected with wildtype *Pf*3D7 parasites were heparin synchronized as described (40) and allowed to reinvade for 4 h before sorbitol synchronization (41) to remove remaining schizonts. Parasites were cultured for a cycle and samples were taken at six hourly intervals and lysed in 0.1% (w/v) saponin. Parasite lysates were separated on 4 - 15% Mini-PROTEAN® TGX™ Gels (Biorad) and transferred to a nitrocellulose membrane for Western blotting. After blocking in 3% (w/v) bovine serum albumin (BSA) in PBS, the membrane was incubated with rabbit anti-M17 (1:1000) and rabbit anti-HSP101 (1:1000;(42)) as a loading control. After washing, the blots were incubated with horseradish peroxidase-conjugated secondary antibodies (1:10,000; Thermo Scientific). Protein bands were detected using the Clarity ECl Western blotting substrate (Biorad) and imaged using a Fujifilm LAS-4000 Luminescent Image Analyzer. ImageJ software (NIH, version 151r) was used to measure the intensity of the bands and GraphPad Prism (V.8.4.2) was used to plot the densitometry. Western blotting to confirm expression of epitope tagged PfA-M17 was performed on asynchronous parasite lysates using rabbit anti-M17 (1:1000) and rabbit anti HSP70 (1:1000).

### Sequential solubility assay

Erythrocytes infected with *Pf*3D7 parasites at trophozoite stage were lysed in 0.05% (w/v) saponin in PBS containing a complete protease inhibitor cocktail (Sigma-Aldrich). Sequential solubilization was performed as previously described (43). Briefly, the parasite pellet was resuspended in a hypotonic lysis buffer (1 mM HEPES, pH 7.4) and incubated on ice for 30 min before undergoing three rounds of freeze-thawing in liquid nitrogen. The solution was then centrifuged at 100,000 *g* for 30 min at 2°C and the supernatant, containing the soluble proteins, collected for analysis. The pellet was then resuspended in 0.1 M Na_2_CO_3_ (pH 11.5), incubated for 30 min and centrifuged, followed by collection of the supernatant fraction, which contained membrane-associated proteins. The remaining pellet was resuspended in 1% (v/v) Triton X-100 in PBS and incubated at room temperature before undergoing centrifugation, with the supernatant containing integral membrane proteins. The final pellet, which contained the insoluble proteins, was resuspended in 1% (v/v) Triton X-100 in PBS. All samples underwent SDS-PAGE gel electrophoresis and Western blotting, with membranes probed with the following antibodies: rabbit anti-M17 (1:1000), rabbit anti-GAPDH (1:1000), rabbit anti-HSP101 (1:1000(42)) and rabbit anti-EXP2 (1:1000(44)).

### Immunofluorescence analysis (IFA)

Transgenic parasites were smeared onto glass slides and allowed to dry overnight before being fixed using acetone:methanol (90:10) for 2 min at -20°C. Slides were air dried before being placed at -20°C until analysis. For immunofluorescent assays, slides were thawed at 37°C for 10 min and blocked with 1 % (w/v) BSA for 1 h. Primary antibody rat anti-HA (Life Technologies) was diluted 1:250 in 0.5% BSA and applied to slides for 2 h before three 5 min washes in PBS. The appropriate AlexaFluor 488-conjugated secondary antibody (1:1000; Life Technologies) was diluted in 0.5% BSA and incubated on slides for 1 h before being washed for 5 min three times in PBS. Cover slips were mounted using Prolong Gold Antifade reagent containing 4’,6-diamidino-2-phenylindole (DAPI; Life Technologies) and incubated overnight at 37°C. Images were taken on a Nikon Eclipse Ti2 microscope at 100 x magnification under oil immersion and processed using ImageJ software (NIH, version 1.53c).

### Knockdown of *Pf*A-M17 expression in *P. falciparum* and growth analysis

Heparin synchronized *Pf*M17-HAglmS parasites were treated with 2.5 mM glucosamine (GlcN) at 0-4 h post-invasion; untreated parasites and *Pf*3D7 parasites treated with 2.5 mM glucosamine served as controls. Parasites were harvested in the first cycle of GlcN treatment (cycle 1) at trophozoite stage, as well as the cycle following (cycle 2) at the same stage. Harvested parasites were lysed in 0.05% saponin, separated by SDS-PAGE electrophoresis and knockdown was analyzed by Western blotting using mouse anti-HA (1:1000; Sigma) antibody with rabbit anti-EXP2 (1:1000) as a loading control. Assessment of parasite growth ± GlcN was determined by analysis of Giemsa-stained smears and images taken with a SC50 5-megapixel or IX71 color camera (Olympus). The parasitemia was determined daily for 6 days following ± GlcN treatment by counting a minimum of 1000 cells. For each day, the stage of development was also determined and plotted on GraphPad Prism (V.8.4.1). Survival of parasites seeded in 96-well plates at 100 parasites per well was determined after 10 days in culture by Sybr Green I assay. Briefly, after freeze-thawing at -80°C, an equal volumes of Lysis buffer (20 mM Tris pH 7.5, 5 mM EDTA, 0.008% saponin (w/v) & 0.008% Triton x-100 (v/v))(45) containing 0.2 µL/mL SYBR Green I Nucleic Acid Gel Stain (10,000x in DMSO; ThermoFisher) was added to each well and incubated for 1 h at RT before fluorescence intensity was read on a Glomax® Explorer Fully Loaded (Promega) with emission wavelengths of 500-550 nm and an excitation wavelength of 475 nm and graphs were generated using GraphPad Prism (V.8.4.1). All experiments were performed in 3 biological replicates and unless otherwise stated, a two-tailed unpaired t-test was used herein to determine statistical significance, with data presented as the mean and error bars representative of standard deviation.

### Hemozoin Assay

Erythrocytes infected with *Pf*M17-HAglmS or *Pf*3D7 underwent sorbitol synchronization before cultures were treated ± GlcN and incubated for a further cycle until parasites developed into trophozoites. Once parasites reached developmentally similar stages, pellets were resuspended in 800 µL of 2.5% SDS in 0.1 M sodium bicarbonate pH 8.8, and then incubated at RT with rotation for 20 min, before undergoing centrifugation at 13,000 *g* for 10 min. Each pellet was then washed twice with 1 mL of the same solution before being resuspending in 500 µL of 5% SDS, 50 mM NaOH and incubating for a further 20 min with rotation. The quantity of monomeric heme was then measured at 405 nM on a Glomax® Explorer Fully Loaded (Promega). Three or four biological replicates were performed, and the data plotted using GraphPad Prism 9 where significance was determined by a two-tailed unpaired t-test. Parasites were assessed for multiple DVs via Geimsa staining and under 100 x magnification with oil, with a combined ≥ 200 parasites counted from 4 biological replicates for each treatment group. A Kruskal-Wallis test with Dunn’s post hoc multiple comparison test was used to determine the significance.

### Determination of compound EC_50_

Parasite viability assays were adapted from previously described methods (46). Briefly, sorbitol synchronized ring stage *Pf*3D7 parasites were cultured in 96-well U-bottom plates at 0.5% parasitemia and 2% hematocrit, to which 50 µL of serially diluted **3,** artemisinin or artesunate (Sigma) was added. After 72 h under standard culturing conditions, plates were placed at -80°C before analysis using the SYBR Green I assay as described above. Uninfected RBCs and parasites treated with the vehicle control DMSO were used to normalize fluorescence. Data from 3 or 4 biological replicates performed in triplicate was plotted as 4-parameter log dose nonlinear regression analysis with a sigmoidal dose-response curve fitted using GraphPad Prism 9 to generate the EC_50_ values, with error bars representative of the SEM.

### Parasite killing rate assay

Assay was performed as previously described with some alterations (47). Briefly, sorbitol synchronized ring or trophozoite stage parasites were cultured in the presence of 10x the EC_50_ of **3** or artesunate for either 24 or 48 h. Cultures that were incubated with either compound for 48 h were fed at 24 h with fresh media containing 10x the EC_50_ of compound. At the completion of these times, RBCs were thoroughly washed to remove the compound and cultures were diluted 1/3 with fresh media and further grown for 48 h before aliquots were placed at -80°C. Cultures were thawed and analyzed using SYBR Green I assay as described above. Parasite viability was determined as a percentage of DMSO-treated parasites cultured alongside compound treated parasites. Artesunate was used as a positive control and for comparison of the parasite killing rate, and experiments were performed in 4 biological replicates.

### Sample preparation for metabolomics analysis

For experiment 1 (Fig. 7) and 2 (Fig. S4), heparin synchronized *Pf*3D7 and *Pf*M17-HAglmS parasites were allowed to invade RBC for 4 h and any remaining schizonts were lysed by sorbitol synchronization. For experiment 1, parasite cultures were then treated for ∼ 36 h with 2.5 mM GlcN or for 1 h with **3** at 10x the EC_50_ (*Pf*3D7 only) or left untreated, while for experiment 2, parasite cultures were only treated with GlcN. Parasites were harvested at developmentally similar timepoints by centrifugation at 900 *g* for 5 min and then resuspended in 10 mL of chilled PBS. Parasite metabolism was quenched by cooling samples to between 3-5°C in an ethanol-dry ice bath. The rest of the preparation was performed at 4°C. Parasites were magnet purified on a VarioMACS column and 3×10^7^ parasites were used for downstream analysis. For experiment 3 (Fig. S5), *Pf*3D7 cultures underwent double sorbitol synchronization 14 h apart, followed by further incubation for 28-42 h to achieve the desired trophozoite stage (28 hpi) at 6% parasitaemia and 2% hematocrit. Infected RBCs (2×10^8^) were treated with 10x the EC_50_ of **3** for 1 h, after which metabolites were extracted. All samples (from experiment 1, 2 and 3) were centrifuged at 650 *g* for 3 min, the supernatant was removed, and the pellet washed in 500 µL of ice-cold PBS. Samples were again centrifuged at 650 *g* for 3 min and pellets were resuspended in 150 µL of ice-cold extraction buffer (100% methanol) and quickly resuspended. The samples were then incubated on a vortex mixer for 1 h at 4°C before being centrifuged at 17,000 *g* for 10 min; from this 100 µL of supernatant was collected and stored at - 80°C until analysis. For each sample, another 10 µL was collected and pooled, to serve as a quality control (QC) sample.

### Liquid chromatography-mass spectrometry (LC-MS) analysis

Liquid chromatography-mass spectrometry (LC-MS) data was acquired on a Q-Exactive Orbitrap mass spectrometer (Thermo Scientific) coupled with high-performance liquid chromatography system (HPLC, Dionex Ultimate® 3000 RS, Thermo Scientific) as described previously (48). Briefly, chromatographic separation was performed on ZIC-pHILIC column equipped with a guard (5 µm, 4.6 × 150 mm, SeQuant®, Merck). The mobile phase (A) was 20 mM ammonium carbonate (Sigma Aldrich), (B) acetonitrile (Burdick and Jackson) and needle wash solution was 50% isopropanol. The column flow rate was maintained at 0.3 mL/min with temperature at 25 °C and the gradient program was: 80% B to 50% B over 15 min, then to 5% B at 18 min until 21 min, increasing to 80% B at 24 min until 32 min. Total run time was 32 min with an injection volume of 10 µL. Mass spectrometer was operated in full scan mode with positive and negative polarity switching at 35,000 resolution at 200 *m*/*z* with detection range of 85 to 1275 *m/z*, AGC target was 1e6 ions with a maximum injection time of 50 ms. Electro-spray ionization source (ESI) was set to 4.0 kV voltage for positive and negative mode, sheath gas was set to 50, aux gas to 20 and sweep gas to 2 arbitrary units, capillary temperature 300 °C, probe heater temperature 120 °C. Approximately 350 authentic metabolite standards were analyzed at the start of each batch to provide accurate retention times to facilitate metabolite identification. Metabolomics samples from each experiment were analyzed as single batches in random order with periodic injections of the pooled QC, and blank samples, to assess analytical quality and aid downstream metabolite identification procedures.

### Metabolomics LC-MS data processing

The acquired LCMS data was processed in untargeted fashion using open source software, IDEOM (49) (http://mzmatch.sourceforge.net/ideom.php) using an updated metabolite database containing all peptides of proteinergic amino acids up to 5 amino acids in length. Initially, *ProteoWizard* was used to convert raw LC-MS files to *mzXML* format and *XCMS* (Centwave) to pick peaks and convert to *peakML* files. *Mzmatch.R* was used for alignment and annotation of related metabolite peaks with a minimum detectable intensity of 100,000, relative standard deviation (RSD) of <0.5 (reproducibility), and peak shape (codadw) of >0.8. Default IDEOM parameters were used to eliminate unwanted noise and artefact peaks. Loss or gain of a proton was corrected in negative and positive ESI mode, respectively, followed by putative identification of metabolites by accurate mass within 3 ppm mass error searching against the IDEOM metabolite database. To reduce the number of false positive identifications, retention time error was calculated for each putative ID using IDEOM built-in retention time model which uses actual retention time data of authentic standards (∼350 standards). Principal-component analysis (PCA) and hierarchical clustering algorithms were performed in Metaboanalyst (50). Metabolomics data are presented as relative abundance values from 3 technical replicates for experiment 1 and 2, while for experiment 3, data is from 4-9 biological replicates. Differences were determined using Welch’s t test where significant interactions were observed. Significance was determined at p values < 0.05. To assess whether identified peptides could be derived from hemoglobin, the sequences of human hemoglobin α, β and δ chains (HBA_HUMAN P69905, HBB_HUMAN P68871, HBD_HUMAN P02042) were searched for any peptide with monoisotopic mass within 0.002 *m/z* of the identified peptide using custom Python scripts.

## Acknowledgments

We thank the Australian Red Cross for providing red blood cells used in this study. We would also like to thank Professor Susan Charman and the Centre for Drug Candidate Optimisation (CDCO) team for providing compound analysis. R.C.S.E. and T.R.M. were recipients of an Australian Government Research Training Program Stipend. This work was supported by an NHMRC Synergy Grant (1185354). T.F.dK.-W is the recipient of an NHMRC Fellowship (1136300).

## Author Contributions

R.C.S.E. conceptualized the study, performed experiments, analyzed data, and wrote the paper. G.S. performed experiments, analyzed data, and wrote the paper. K.H created the transgenic parasite line. N.A.C performed experiments, analyzed data, supervised work, and wrote the paper. T.R.M. performed experiments and C.T.W. analyzed data. P.J.S. designed the compound, supervised work, and wrote the paper. N.B.V. synthesized the compound and contributed to its design. C.A.M. analyzed data and wrote the paper. D.C. supervised work and wrote the paper. S.M. conceptualized the study, performed experiments, analyzed data, supervised work, and wrote the paper. T.F.dK.-W conceptualized the study, performed experiments, analyzed data, supervised work and wrote the paper. All authors edited the final paper.

