## Supplementary Data for "Genetic and chemical validation of *Plasmodium falciparum* aminopeptidase *Pf*A-M17 as a drug target in the hemoglobin digestion pathway"

### Supplementary text: Structure-activity relationship of 3, a specific PfA-M17 inhibitor.

The design of **3** largely focused on reducing the polarity and improving water solubility relative to the current front-runner compound of our hydroxamic-acid based inhibitor series, *N*-(2-(hydroxyamino)-2-oxo-1-(3′,4′,5′-trifluoro-[1,1′-biphenyl]-4-yl)ethyl)-3,3-dimethylbutanamide (**6l** in [19]). This was addressed by replacing the 3,4,5-trifluorophenyl moiety present in **6l** with a 4-hydroxymethylphenyl. This change reduced the cLogP and provided a hydrogen bond donor which proved successful in significantly improving water solubility (Table 1 below). Metabolic stability, as assessed with an *in vitro* liver microsome assay, was also improved.

**Table 1. Polarity, solubility and metabolic stability of 3**

| Compound | cLogP[a] | Solubility[b] (mg/mL) | Metabolic stability[c] ($t_{½}$, min) |
|---|---|---|---|
| **6l** from ref 19 | 3.6 | 12.5-25 | 149 |
| **3** | 2.5 | >100 | >255 |

[a] Calculated using the ChemAxon chemistry cartridge via JChem for Excel software (version 16.4.11).

[b] Compound in DMSO was spiked into pH 6.5 phosphate buffer with the final DMSO concentration being 1%. After 30 min had elapsed, samples were then analyzed via Nephelometry to determine a solubility range.

[c] The metabolic stability assay was performed by incubating the test compound in liver microsomes at 37°C and a protein concentration of 0.4 mg/mL. The metabolic reaction was initiated by the addition of an NADPH-regenerating system and quenched at various time points over a 60 min incubation period by the addition of acetonitrile containing diazepam as internal standard. Control samples (containing no NADPH) were included (and quenched at 2, 30 and 60 min) to monitor for potential degradation in the absence of cofactor.

Aminopeptidase activity assays clearly show that **3** is a selective and potent inhibitor of *Pf*A-M17. To confirm the binding mode and suggest a reason for such excellent selectivity, we solved the X-ray crystal structure of *Pf*A-M17 bound to **3**. Crystals of *Pf*A-M17 were presoaked in solution containing excess inhibitor prior to data collection whereby the crystals diffracted to a modest 2.49 Å. Molecular replacement using the unbound X-ray crystal structure of *Pf*A-M17 (PDB ID 3KQZ) showed clear Fo-Fc density for the ligand in each active site of the two complete *Pf*A-M17 hexamers that were present in the asymmetric unit. The inhibitor binding modes were well conserved across all twelve active sites and subsequently from here on in the molecular interactions of **3** with the active site is described only for Chain A.

Similar to our previous structural data from our hydroxamic core, we observed that the position of the hydroxamic acid (zinc binding group), pivalamide (S1' pocket substituent) and the biphenyl ring system (S1 pocket substituent) were conserved. The two zinc ions are coordinated by the hydroxamic oxygens, that also form hydrogen bonds with the conserved carbonate ion and catalytic residue Lys386 (Supplementary Fig. 3). In the S1 pocket, the *tert*-butyl makes no interactions with the body of the protein, but the carbonyl 3 forms a hydrogen bond with the main chain amine of Gly489 as well as a water molecule (Supplementary Fig. 3). The 4-hydroxymethylphenyl group that replaced the trifluorophenyl group present in **6f**[19] packed with the hydrophobic residues Leu487, Gly489, Leu492 Met396, Phe583 and Ala577 and the hydroxyl group of **3** can interact with the sulfur atom of Met392 (Fig. 4B, Supplementary Fig. 3).

To try and understand why **3** could act as a selective inhibitor of *Pf*A-M17 and showed very little activity toward *Pf*A-M1, we attempted to solve the X-ray crystal structure of *Pf*A-M1 bound to **3**. This was unsuccessful and no compound density was observed within any datasets collected from co-crystalized or soaked *Pf*A-M1 crystals. This was not surprising as our previous attempts to collect structures of *Pf*A-M1 crystals bound with weak inhibitors have always failed. Using the structure 4ZX4.pdb as a template, we were able to superpose **3** onto the 4ZX4.pdb ligand however inspection of the S1 pocket with the superposed ligand does not identify why this compound shows low potency toward *Pf*A-M1. Replacement of the trifluorophenyl group for 4-hydroxymethylphenyl positioned the hydroxyl group of **3** close to E572 in the S1 pocket (~ 2.8 Å from the Cα atom and 1.7 Å from the Cγ atom). However, it would be surprising if this close contact was the reason for the lack of potency for this inhibitor in that E572 has been shown to move to accommodate bulky hydrophobic groups that extend from longer inhibitors[50] and the hydroxyl group and the E572 side-chain would likely be able to re-position/rotate to avoid a clash. In the S1' pocket, the pivalamide moiety is easily positioned to the same place as other compounds with a similar scaffold but also does not suggest an obvious reason for the lack of potency with regard to the fit of the compound into the active site or its substrate pockets. It may be that the change in electronegativity, via the loss of the 3 fluorine atoms, results in the compound not being able to access the buried active site with the same affinity.

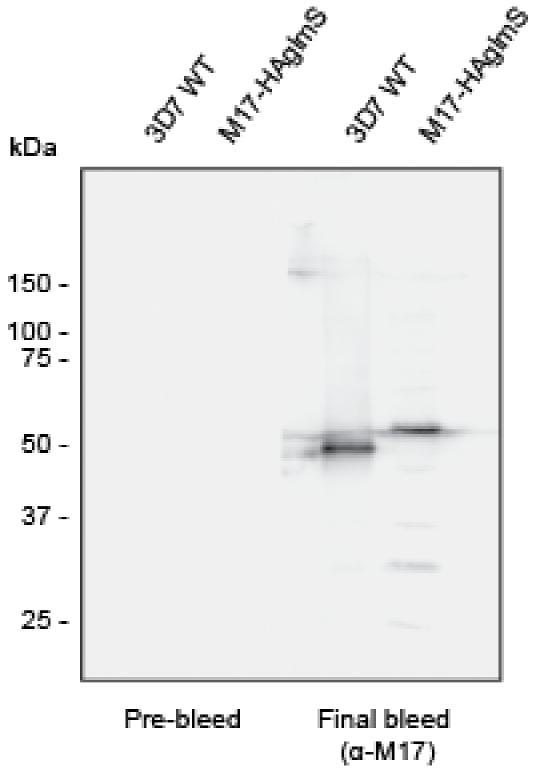


**Supplementary Figure 1.** Western blot of lysates prepared from mixed stage *Pf*3D7 wild type (WT) and *Pf*A-M17-HAglmS parasites probed with either pre-bleed rabbit serum or rabbit serum after multiple rounds of inoculation with *Pf*A-M17 recombinant protein (final bleed). The expected molecular mass of *Pf*A-M17 is 68 kDa.

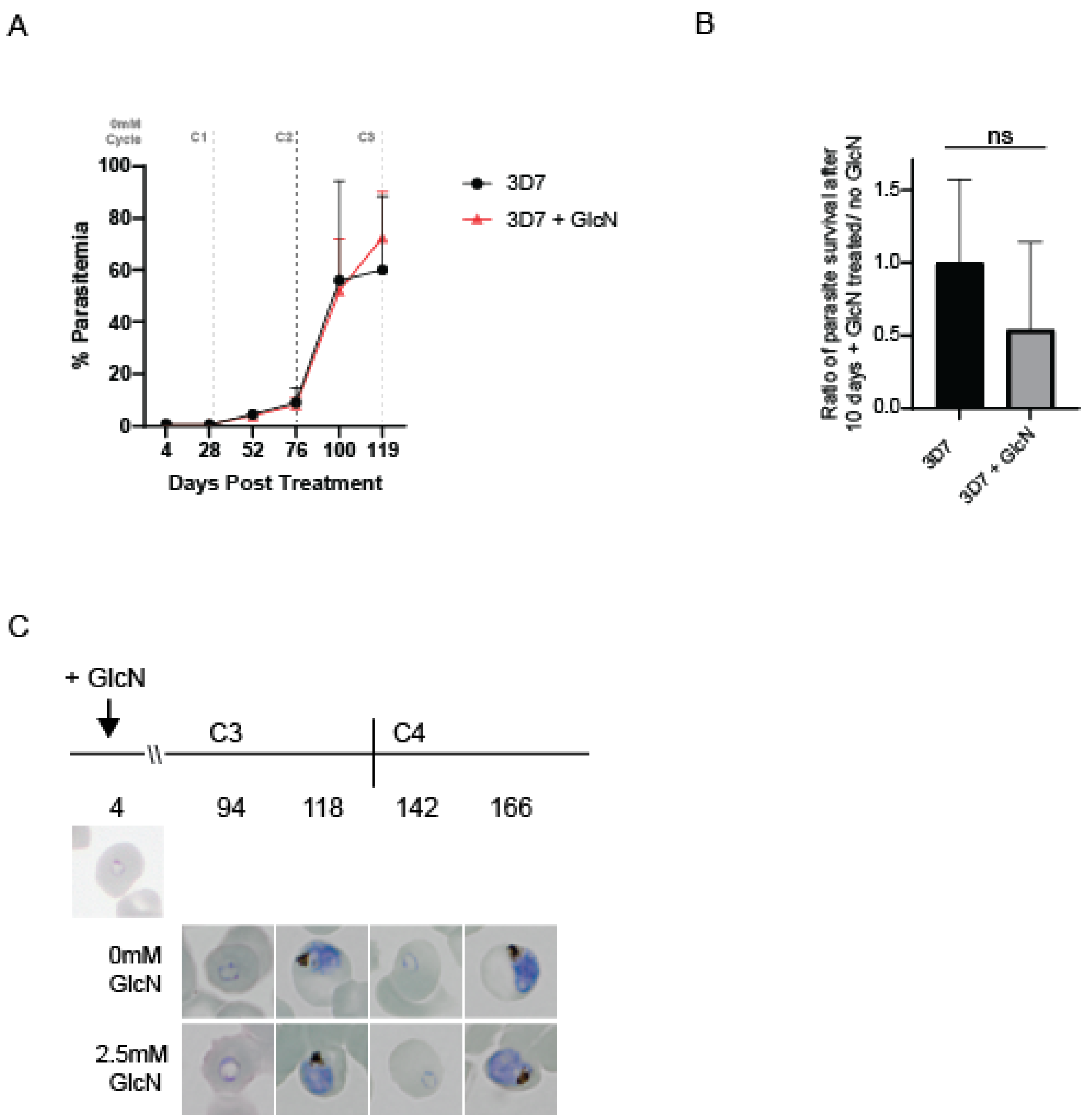


**Supplementary Figure 2.** Addition of Glucosamine does not significantly affect growth of *Pf*3D7 parasites. **(A)** Parasitemias of *Pf*3D7 parasites cultured ± GlcN over 3 cycles shows that parasite growth is not significantly affected between treatment groups. Shown is the mean ± standard deviation (n= 2). **(B)** Ratio of parasite survival of *Pf*3D7 after treatment with GlcN for 10 days compared to untreated parasites as determined by Sybr Green 1 assay. Shown is the mean ± standard deviation (n= 2). Statistical significance was determined using an unpaired t-test. **(C)** Representative Giemsa-stained parasite smears of *Pf*3D7 cultures treated with 0 mM or 2.5 mM GlcN.

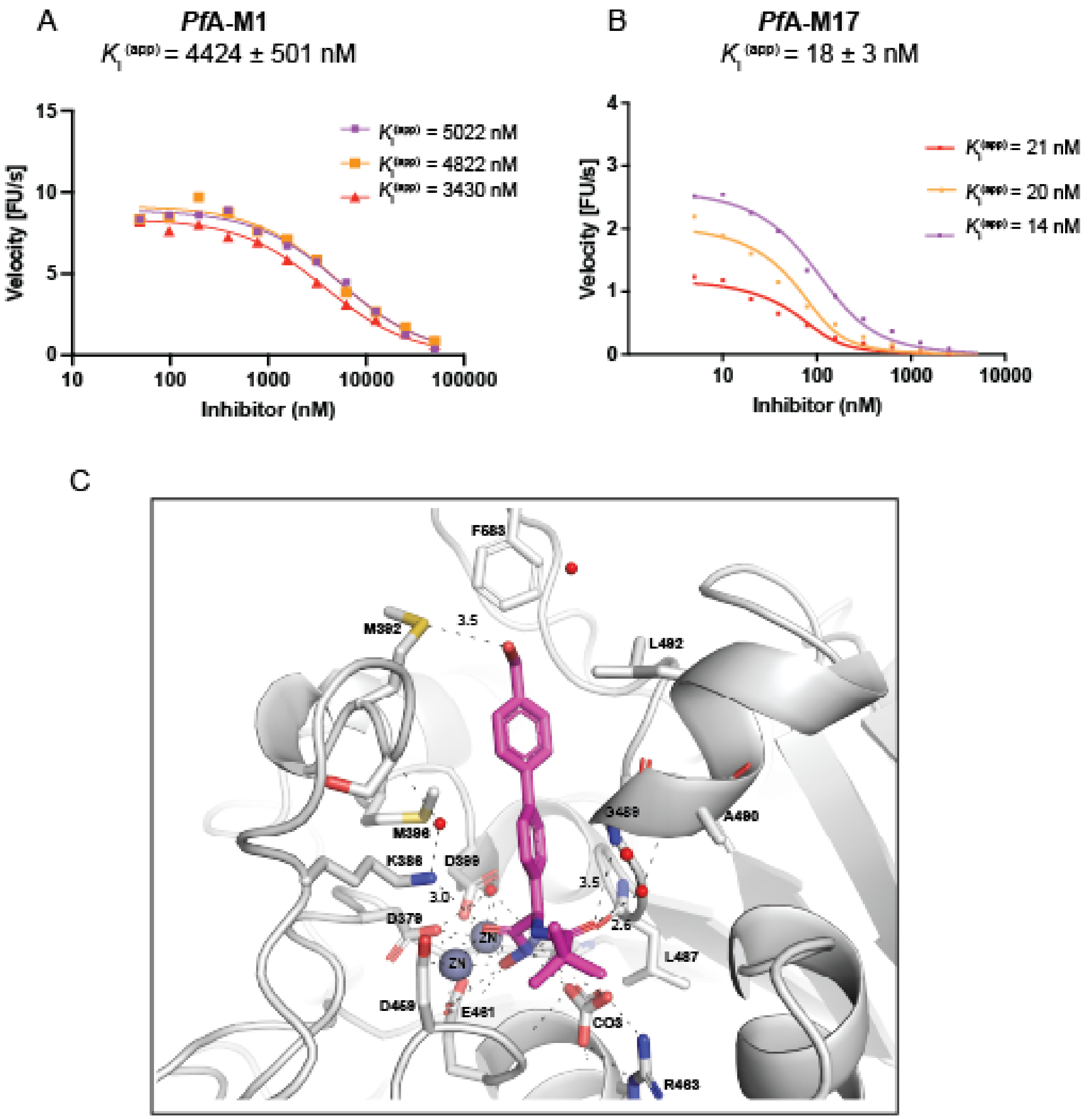


**Supplementary Figure 3:** Compound **3** is a potent and selective *Pf*A-M17 inhibitor. Dose response curves showing aminopeptidase activity (fluorescence units per sec, FU/s) in the presence of increasing concentration (shown as log nM) of **3** for *Pf*A-M1 (**A**) and *Pf*A-M17 (**B**). Three separate dose-response curves prepared from three separate protein purifications are shown, with the final $K_i^{(app)}$ value is the mean ± SEM of the three independent values (n=3).

(**C**) Binding of **3** (magenta sticks) to *Pf*A-M17 (grey cartoon). Interactions between **3** and *Pf*A-M17 are shown by black dash lines and distances (Å) of key interactions (excluding zinc coordination) are indicated above dashed lines. Residues involved in hydrogen bonding and hydrophobic packing interactions are shown in grey sticks and residues numbers are indicated.

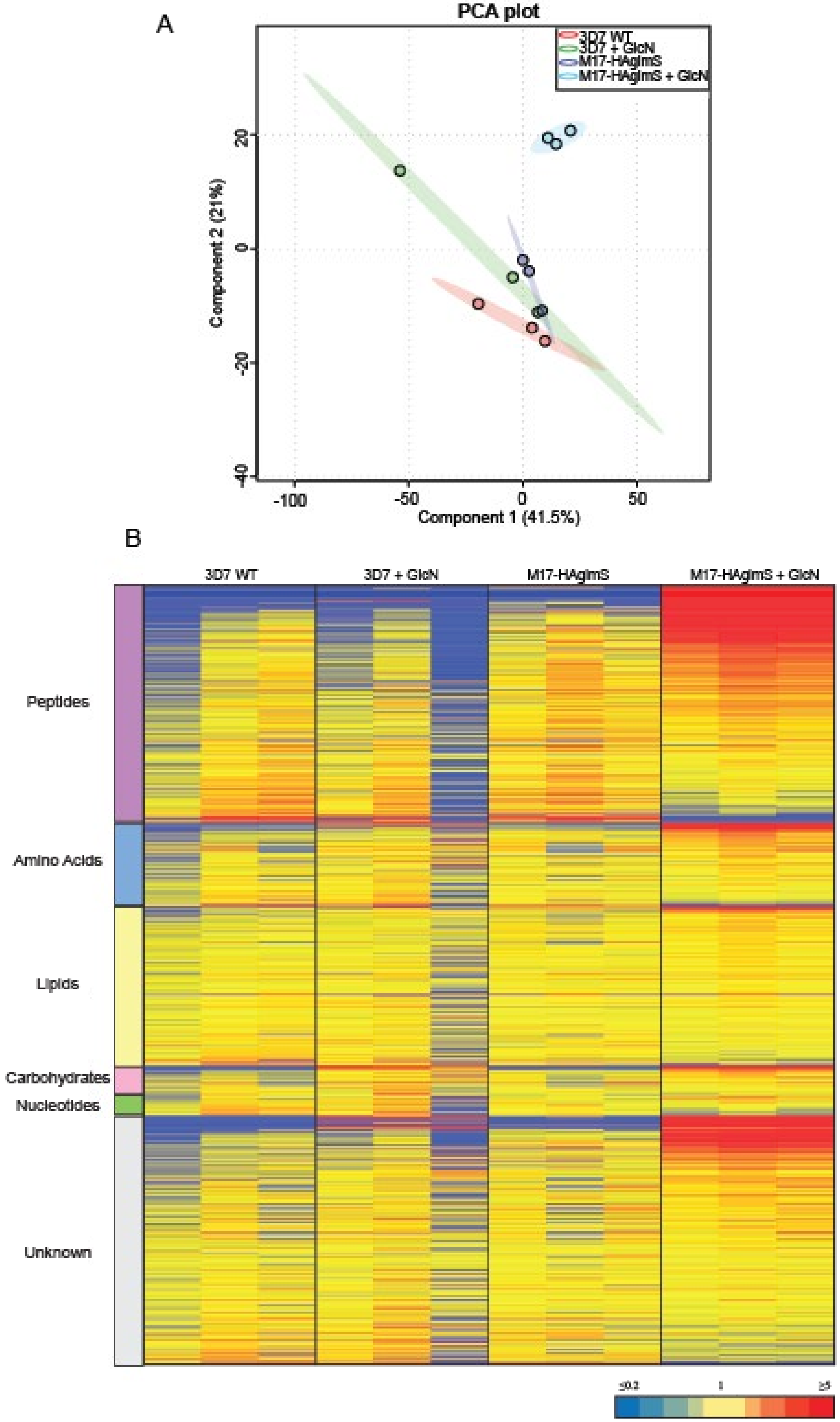
A
PCA plot
3D7 WT
3D7 + GlcN
M17-HAglmS
M17-HAglmS + GlcN
Component 2 (21%)
Component 1 (41.5%)
20
0
-20
-40
-100
-50
0
50
B
3D7 WT
3D7 + GlcN
M17-HAglmS
M17-HAglmS + GlcN
Peptides
Amino Acids
Lipids
Carbohydrates
Nucleotides
Unknown
≤0.2
1
≥5

**Supplementary Figure 4.** Untargeted metabolomics analysis of *Pf*A-M17-HAglmS and *Pf*3D7 parasites treated with ± GlcN from experiment 2. (**A**) Principal component analysis (PCA) of parasites (*Pf*A-M17-HAglmS and *Pf*3D7) treated with +/- GlcN. Scores plot show principal components one and two, data points indicate individual sample replicates within each condition and the shaded area denotes 95% confidence interval. (**B**) Heatmap analysis of peak intensities of all putative metabolites for each condition. Data is shown from three technical replicates, red, blue and yellow indicates increase, decrease and no change respectively in the relative abundance of putative metabolites identified.

A

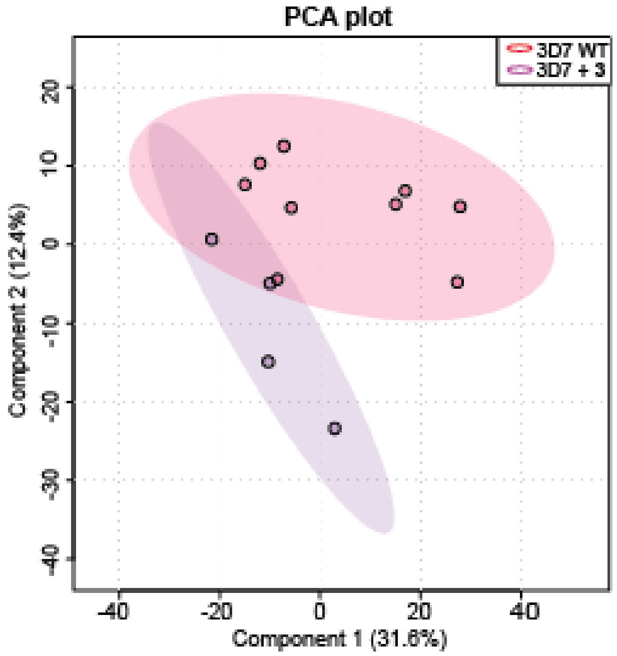
PCA plot
3D7 WT
3D7 + 3
Component 2 (12.4%)
Component 1 (31.6%)
-40
-20
0
20
40
20
10
0
-10
-20
-30
-40

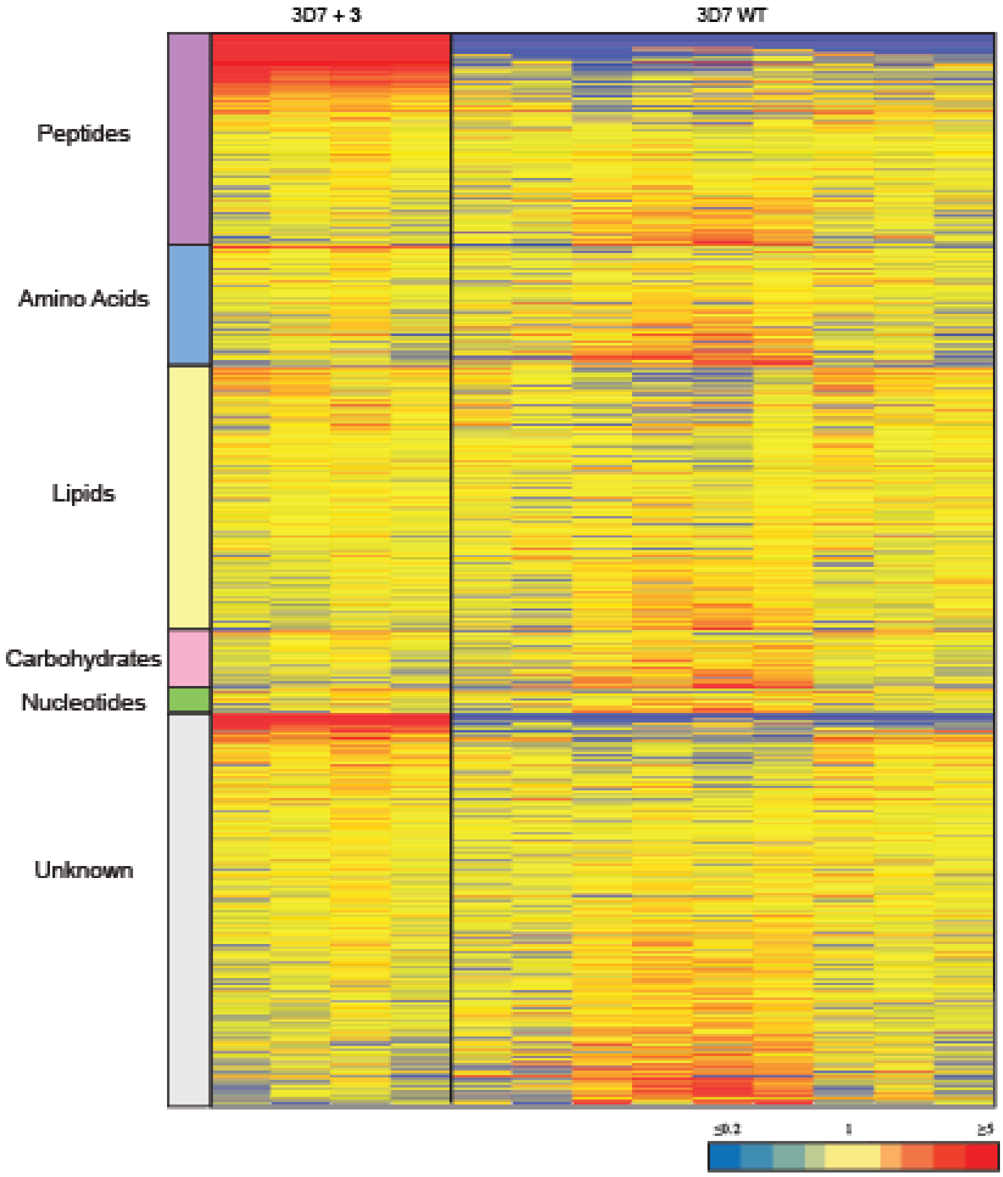
3D7 + 3
3D7 WT
Peptides
Amino Acids
Lipids
Carbohydrates
Nucleotides
Unknown
≤0.2
1
≥5

**Supplementary Figure 5.** Untargeted metabolomics analysis of *Pf*3D7 parasites treated with **3** and DMSO control from experiment 3. (**A**) Principal component analysis (PCA) of parasites *Pf*3D7 treated with **3** and DMSO control. Scores plot show principal components one and two, data points indicate individual sample replicates within each condition and the shaded area denotes 95% confidence interval. (**B**) Heatmap analysis of peak intensities of all putative metabolites for each condition. Data is shown from four-nine biological replicates, red, blue and yellow indicates increase, decrease and no change respectively in the relative abundance of putative metabolites identified.

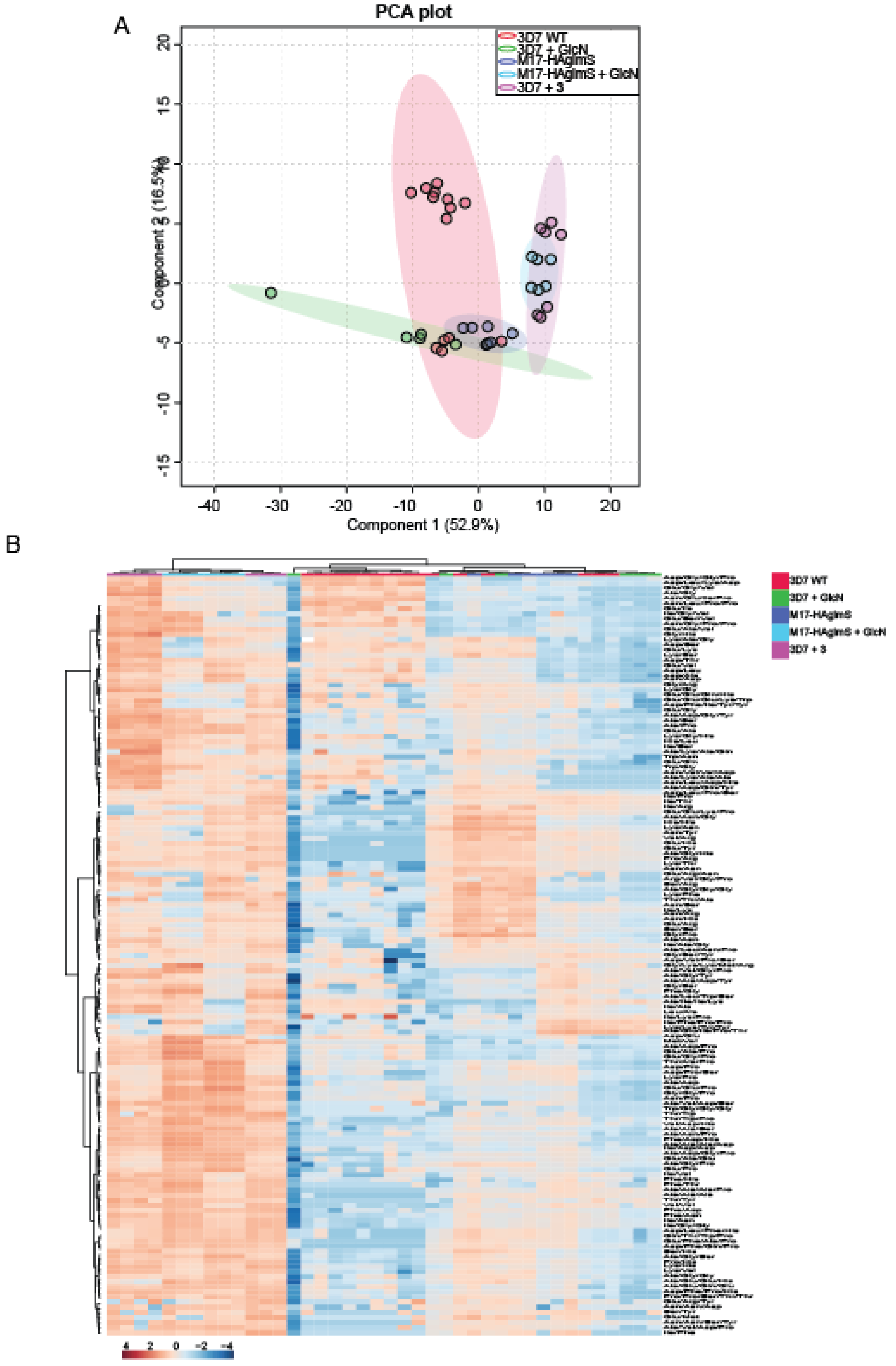
A
PCA plot
3D7 WT
3D7 + GlcN
M17-HAglmS
M17-HAglmS + GlcN
3D7 + 3
Component 2 (16.5%)
Component 1 (52.9%)
B
3D7 WT
3D7 + GlcN
M17-HAglmS
M17-HAglmS + GlcN
3D7 + 3

**Supplementary Figure 6.** Targeted analysis of all common peptides identified from experiment 1, 2 and 3. (**A**) Principal component analysis (PCA) of common peptides identified across three experiments for parasites (*Pf*A-M17-HAglmS and *Pf*3D7) treated with +/- GlcN and **3** or DMSO control. Scores plot show principal components one and two, data points indicate individual sample replicates within each condition and the shaded area denotes 95% confidence interval. (**B**) Hierarchical clustering of the common peptides identified across the three independent experiments. Vertical clustering displays similarities between sample groups, while horizontal clusters reveal the relative abundances (median normalised) of common identified peptides (149). The colour scale bar represents $\log_2$ (mean-centred and divided by the standard deviation of each variable) intensity values.

**Supplementary Table 1. Crystallography and refinement statistic for PfA-M17 bound to 3**

| | *Pf*A-M17-**3** |
|---|---|
| PDB ID | 7RIE |
| *Data Collection Statistics* | |
| Wavelength | 0.953727 |
| Resolution range | 48.73 - 2.49 (2.58 - 2.49) |
| Space group | P 21 21 21 |
| Unit cell (a, b, c, α, β, γ) | 175.29 175.86 234.68 90 90 90 |
| Total reflections | 498247 (47143) |
| Unique reflections | 250277 (23847) |
| Multiplicity | 2.0 (2.0) |
| Completeness (%) | 99.45 (95.51) |
| Mean I/sigma(I) | 7.02 (1.30) |
| Wilson B-factor | 41.46 |
| R-pim | 0.06067 (0.3973) |
| CC1/2 | 0.996 (0.727) |
| *Refinement statistics* | |
| Reflections used in refinement | 250101 (23835) |
| Reflections used for R-free | 12429 (1229) |
| R-work | 0.1947 (0.2850) |
| R-free | 0.2281 (0.3315) |
| Number of non-hydrogen atoms | 0.964 (0.818) |
| macromolecules | 0.952 (0.737) |
| ligands | 49071 |
| solvent | 47016 |
| Protein residues | 528 |
| RMS(bonds) | 1527 |
| RMS(angles) | 6182 |
| Ramachandran favored (%) | 0.003 |
| Ramachandran allowed (%) | 0.57 |
| Ramachandran outliers (%) | 97.07 |
| Rotamer outliers (%) | 2.78 |
| Clashscore | 0.15 |
| Average B-factor | 1.17 |
| macromolecules | 2.99 |
| ligands | 43.18 |
| solvent | 43.34 |
| Number of TLS groups | 41.48 |

Statistics for the highest-resolution shell are shown in parentheses.

**Supplementary Table 2. Oligonucleotide sequences used in this study**

| Name | Target | Oligonucleotide Sequence |
|---|---|---|
| DO276 | Ribozyme | GTGATTTCTCTTTGTTCAAGGA |
| DO657 | M17 | AATAACCGCGGGGATCAGTTGCTGATTTAAGT |
| DO658 | M17 | AATAAGGCGCGCCCTAGAGCGTCATTGAGTACAA |
| DO733 | M17 | GACCGATGAAAGGTTCAAATCT |
| DO734 | M17 | GTGAGGTACACCACACATG |
